# Europe PMC Annotated Full-text Corpus for Gene/Proteins, Diseases and Organisms

**DOI:** 10.1101/2023.02.20.529292

**Authors:** Xiao Yang, Shyamasree Saha, Aravind Venkatesan, Santosh Tirunagari, Vid Vartak, Johanna McEntyre

## Abstract

Named entity recognition (NER) is a widely used text-mining and natural language processing (NLP) sub-task. In recent years, deep learning methods have superseded traditional dictionary, and rule-based NER approaches. A high-quality dataset is essential to take full advantage of the recent deep learning advancements. While several gold standard corpora for biomedical entities in abstracts exist, only a few are based on full-text research articles. The Europe PMC literature database routinely annotates Gene/Proteins, Diseases and Organisms entities; to transition this pipeline from a dictionary-based to a machine learning-based approach, we have developed a human-annotated full-text corpus for these entities comprising 300 full-text open access research articles. Over 72,000 mentions of biomedical concepts have been identified within approximately 114,000 sentences. This article describes the corpus and details how to access and reuse this open community resource.

## Background & Summary

Europe PubMed Central (Europe PMC)^1^ is a repository of life science research articles, including peer-reviewed full-text research articles and abstracts, and preprints -all freely available for use via the website (https://europepmc.org). Europe PMC contains over 33.3 million abstracts and 8.7 million full-text articles, and since 2020 adds over 1.7 million new articles yearly. The rapid growth in the number of publications within the biological research space makes tracking research trends and assimilating knowledge a challenging and time-consuming effort. With the digitisation of large portions of biological literature and advancements in natural language processing (NLP)/machine learning (ML), it is now possible to build sophisticated tools and the required infrastructure to process research articles in order to extract biological entities, concepts and relationships in a scalable manner.

Harnessing the NLP techniques, tools such as LitSuggest^2^ and PubTator^3^ are being used in biomedical literature curation^4,5^, recommending relevant biomedical literature, or automatically annotating biomedical concepts^6^, such as genes and mutations, in PubMed abstracts and PubMed Central (PMC) full-text articles. Furthermore, in a step towards FAIRification^7–9^ and sharing text-mined outputs across the scientific community, Europe PMC has established a community platform to capitalise on the advances made. Annotations from various text-mining groups are consolidated and made available via open APIs and a web application called SciLite^10^, which highlights the annotations on the Europe PMC’s website. Several other biological resources including STRING^11^ and neXtProt^12^ have embedded NLP processes in their data workflows to serve their user community better. Developement of such NLP tools require the availability of open data (full-text corpora). Thanks to the biomedical text mining community, which has endorsed open data; resources such as PubMed, PubMed Central and Europe PMC provide open access abstracts and full-text for researchers to download. The COVID-19 Open Research Dataset Challenge (CORD-19 dataset)^13^ is a recent example of using text-mining to tackle specific scientific questions. This dataset consists of full-text scientific articles about COVID-19 and related coronaviruses. Additionally, BioC^14^ provides a subset of those full-text articles in a simple BioC format, which can reduce the efforts of text processing. Biomedical datasets, such as those from BioASQ^15^ and BioNLP^16^ shared tasks, enable the development and testing of novel ideas, including deep learning methodologies. With the development of such biomedical datasets, great improvements in biomedical text mining systems have been made. From the results of recent BioASQ challenges (2013 to 2019), the performance of cutting edge systems keep advancing for tasks such as large-scale semantic indexing and question answering (QA)^17^. while corpora without annotations are good for learning semantics, text-mining tools trained on human-annotated corpora outperform those trained on non-annotated ones. Therefore, open-source gold-standard datasets are crucial for improving biomedical text mining systems. In particular, transformer based deep learning models, such as BERT^18^ and GPT^19^, have show that pre-training language models with large text corpora improves performance on downstream applications. However, compared to the text corpora, gold-standard biomedical datasets with human annotations are expensive to obtain, because they require domain experts to spend significant amounts of time creating accurate annotations. Therefore, generating human-annotated biomedical datasets is valuable for biomedical text mining, because once they are available, machine learning algorithms have an accurate starting point to learn from.

The Colorado Richl Annotated Full-Text Corpus (CRAFT)^20^ is one such human-annotated biomedical dataset, that has been widely applied by researchers to develop and evaluate novel text mining algorithms. It contains 97 full-text open access biomedical journal articles that have both semantic and syntactic annotations, including coreference annotations and 10 biomedical concepts. Besides the CRAFT corpus, other corpora such as BioCreative V CDR corpus (BC5CDR)^21^, BC2GM^22^, Linnaeus^23^, S800^24^, GAD^25^, EUADR^26^, and BioASQ^15^ have also been used to extensively test new algorithms on the concept recognition task. Comprehensive and high-quality annotations make the CRAFT corpus one of the most important gold-standard datasets in the biomedical domain. Recent publications^27,28^ have shown that sophisticated systems can be developed by using annotated biomedical datasets. One example is OGER^27^, a system trained with neural networks that outperforms the previous dictionary-based systems in the concept recognition task. In particular, when evaluated on the CRAFT corpus^20^, more pronounced improvements have been achieved in identifying Chemical, Organism, Protein, and Sequences Concepts^27^. Notably, as pre-trained models such as BERT^18^ have gained traction in the biomedical domain, many systems have been developed by training models with multiple biomedical datasets. For example, the BioBert model^29^ has been trained and evaluated on multiple datasets for downstream tasks including Named Entity Recognition, Relation Extraction and Question Answering.

This study presents the Europe PMC Annotated Full-text Corpus (EPMCA), a collection of 300 research articles from the Europe PMC Open Access subset. The selected articles have been human annotated to indicate mentions of three biomedical concepts; Gene/Protein, Disease, and Organism. Since all annotations are created based on guidelines, this helped the human annotators select the correct text span and type of annotation. Three additional articles that were used in a pilot study are also published with this study. The size of the EPMCA is among the largest human-annotated biomedical corpora. We believe that the high-quality gold standard annotations of the EPMCA corpus will be an important addition to other existing datasets and provide significant benefits for biomedical text mining.

## Methods

The overall strategy for Full-text annotation workflow is presented in Figure 1. Out of a million Open Access (OA) full-text articles archived on the 31st of August 2018 in Europe PMC, a subset of 300 articles were selected as the gold standard for the curation. This section presents the methods we employed to stratify those articles and select the representative gold standard set followed by the annotation guidelines and article annotation.

**Figure 1.**
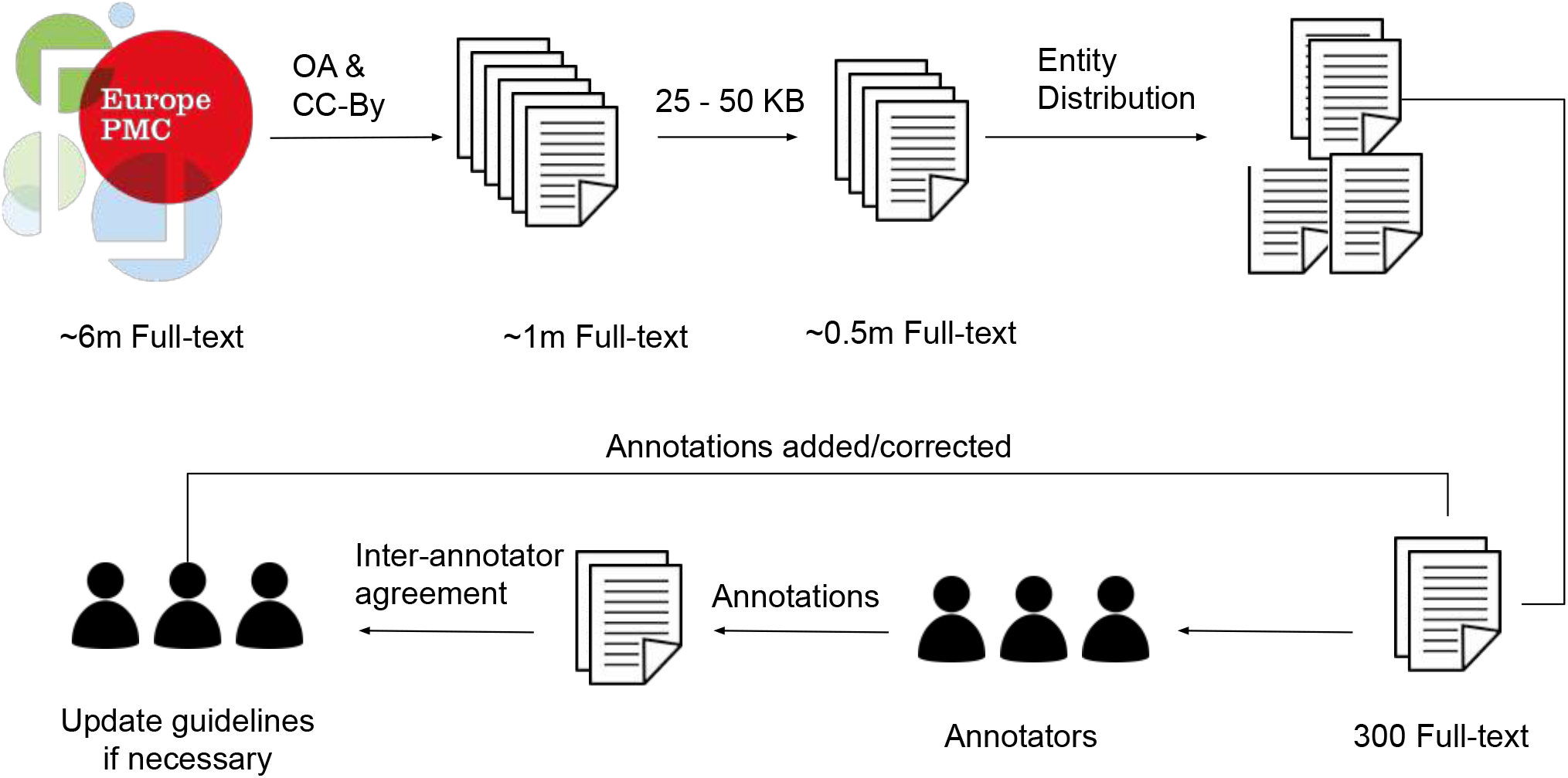
The illustration of the full-text annotation workflow. There were approximately six million full-text articles in the Europe PMC repository archived on the 31st of August, 2018 (v2018.09) of which approximately one million were Open Access (OA) with a CC-BY licence. Thereafter, to have articles specific to research, size between 25 and 50KB were selected, which resulted in a collection of approximately 0.5 million articles. This was followed by sorting the articles with the entity mentions into low, medium, and high bins for each entity type, i.e. Gene/Protein, Disease, and Organisms. Finally, 300 articles were selected that represented the aforementioned entity types for each article. The workflow included working with the annotators iteratively to improve the annotation guidelines.

### The Open Access article set in Europe PMC and CC-BY-licenced articles

Because a primary outcome of this work was to create a training set for anyone to use, the first constraint applied was to use Open Access articles that have a parsable/machine-readable (available in JATS XML standard, for more information please refer to https://jats.nlm.nih.gov/) *CC-BY* licence. We used the archived open access set from 31st August 2018 (v.2018.09) [Available at http://europepmc.org/ftp/archive/] as a basis, which consists of 2, 113,557 articles, of which 991,529 articles had a parsable *CC-BY* licence.

#### Body size

Using the 991,529 *CC-BY* articles as a starting point, we measured the size of the full-text article *<BODY>* section and grouped them into bins of 10*KB* size to find the most representative articles. More than 50% of articles were in the range of 25−50*KB* (Figure 2). Using this size range further constrained the pool to 503,950 articles. Constraining the article size range also meant that the annotators would be provided with a more consistent article set as presumably articles falling outside this range are likely to not be research articles.

**Figure 2.**
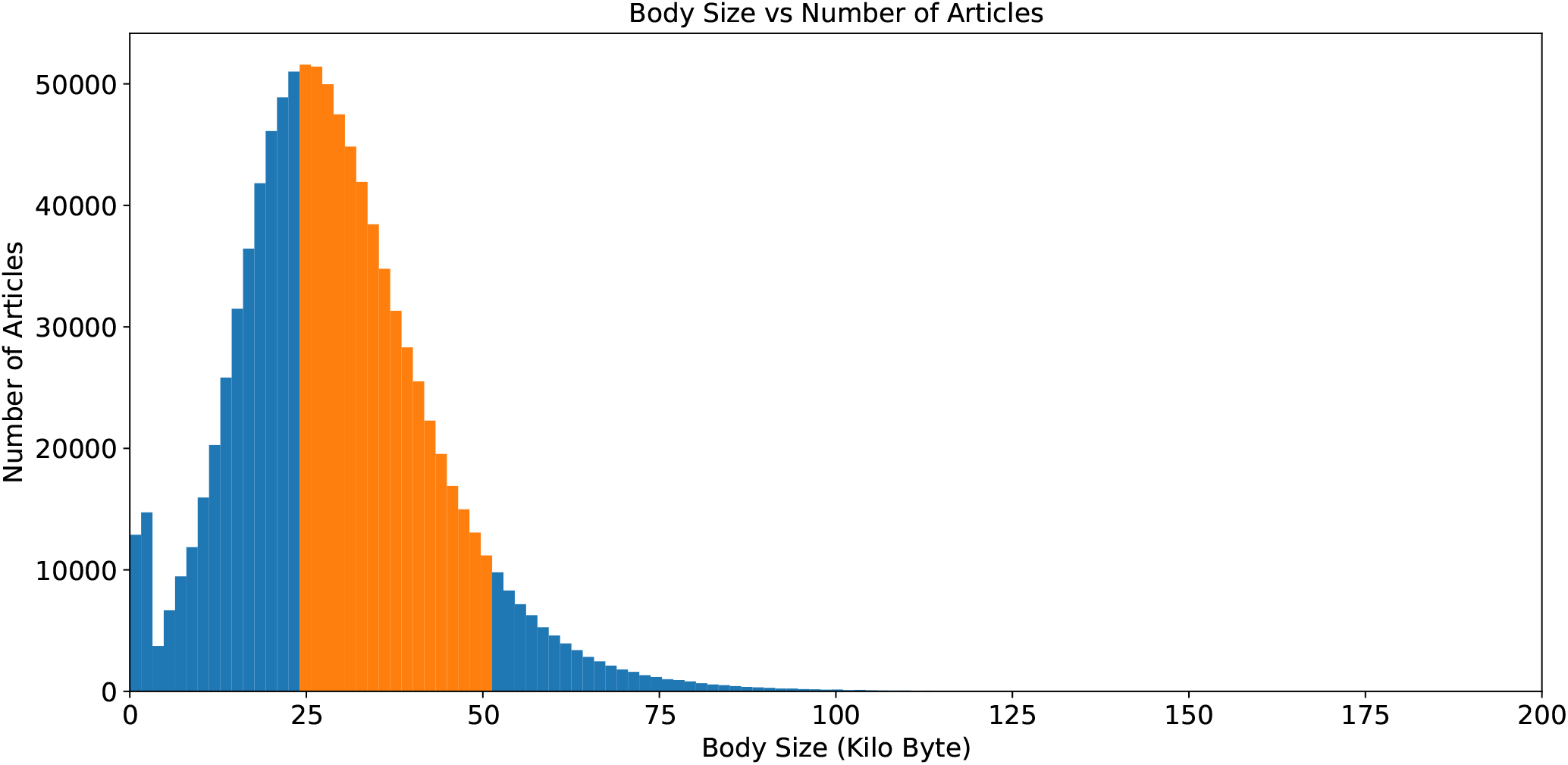
Distribution of body sizes of full-text articles with a CC-BY licence on the 31st August 2018 (v.2018.09) frozen set.

### Entity frequency distribution

The pool of 503,950 “standard-sized” articles were further stratified based on the term frequency of the three entities of interest, namely; Gene/Proteins, Diseases, and Organisms. Using the current Europe PMC dictionary-based annotation pipeline to annotate the articles, we established the range of entity frequencies in the articles (Figure 3) and created high (H), medium (M), and low (L) frequency tertiles by splitting them at the 33 and the 66 percentiles (Table 1). This resulted in 27 bins of articles from these tertiles of three entities (33) (Figure 4). All the articles in the Low-Low-Low bin contain a small number or no mentions of any of the entities but represent the largest number of articles (42,261 articles, more than 8% of total articles). Because these would add little value to the training dataset, this bin was excluded from the article selection process. There were 46,1689 articles in the remaining 26 bins. We then randomly selected 300 articles in total across all 26 bins in proportion to the number of articles in each bin (2–20 articles from a bin in real terms, Figure 5). For example, only two articles were selected from the Low Disease, High Gene/Protein, Low Organism bin.

**Table 1.** The abundance of key entities is used to establish tertile boundaries.

| Entity | Low frequency of occurrence count (L) |  | Medium frequency of occurrence count (M) |  | High frequency of occurrence count (H) |  |
| --- | --- | --- | --- | --- | --- | --- |
|  | Lower | Upper | Lower | Upper | Lower | Upper |
| Genes/Proteins | 0 | 11 | 12 | 80 | 81 | 2408 |
| Organisms | 0 | 9 | 10 | 57 | 58 | 3108 |
| Diseases | 0 | 4 | 5 | 32 | 33 | 678 |

**Figure 3.**
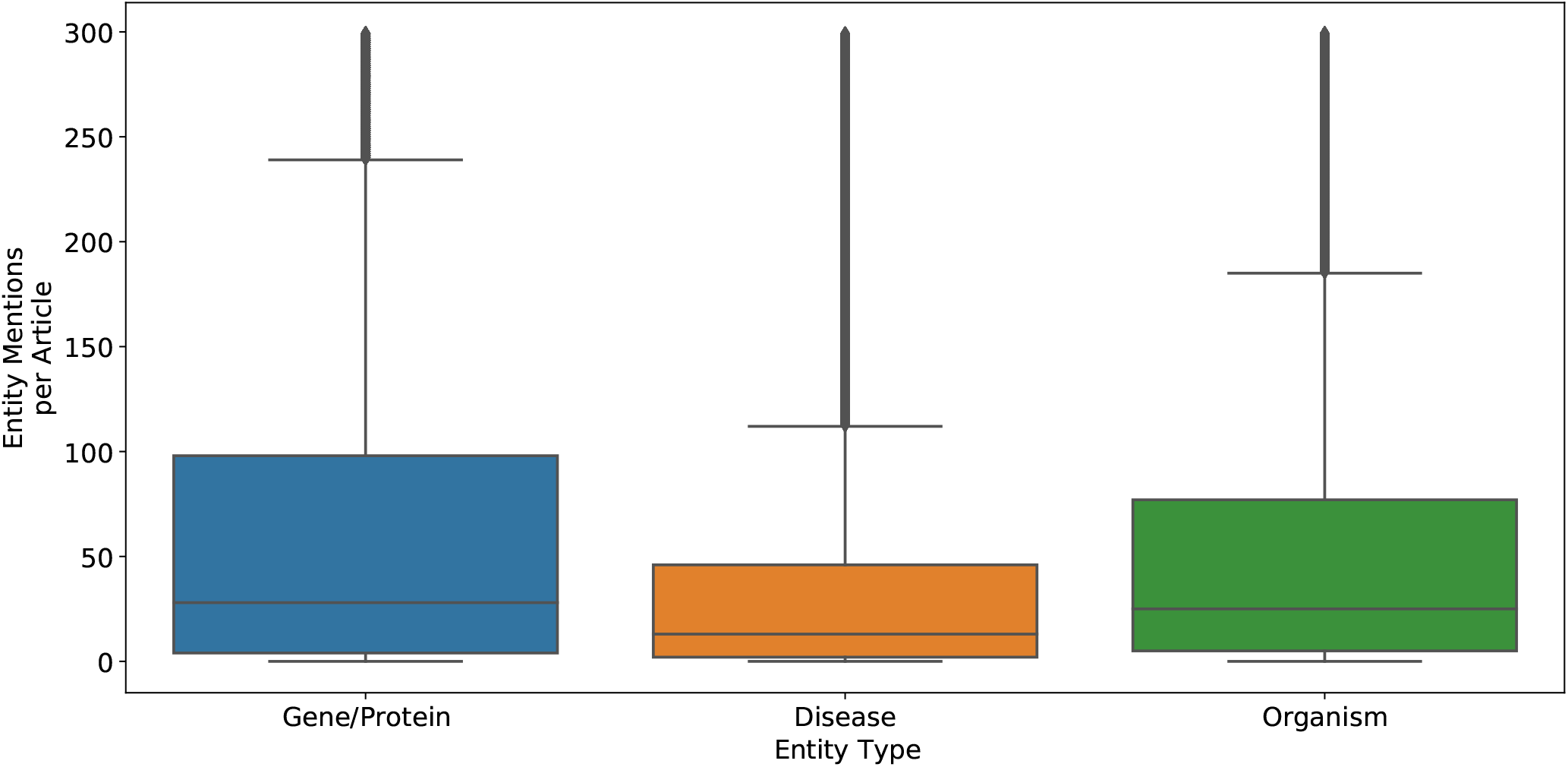
Distribution of entity mentions (Gene, Disease and Organism) per full-text article from the candidate pool. For the convenience of the display, we have used a threshold of a maximum of 300 mentions per article per entity type for this figure, although the maximum was 2408 for Gene/Protein, 678 for Disease, and 3108 for Organism. This figure shows that, on average, Disease mentions are almost half of Gene/Protein mentions per article. This distribution helped us to set entity count boundaries for the article stratification required to select the final corpus.

**Figure 4.**
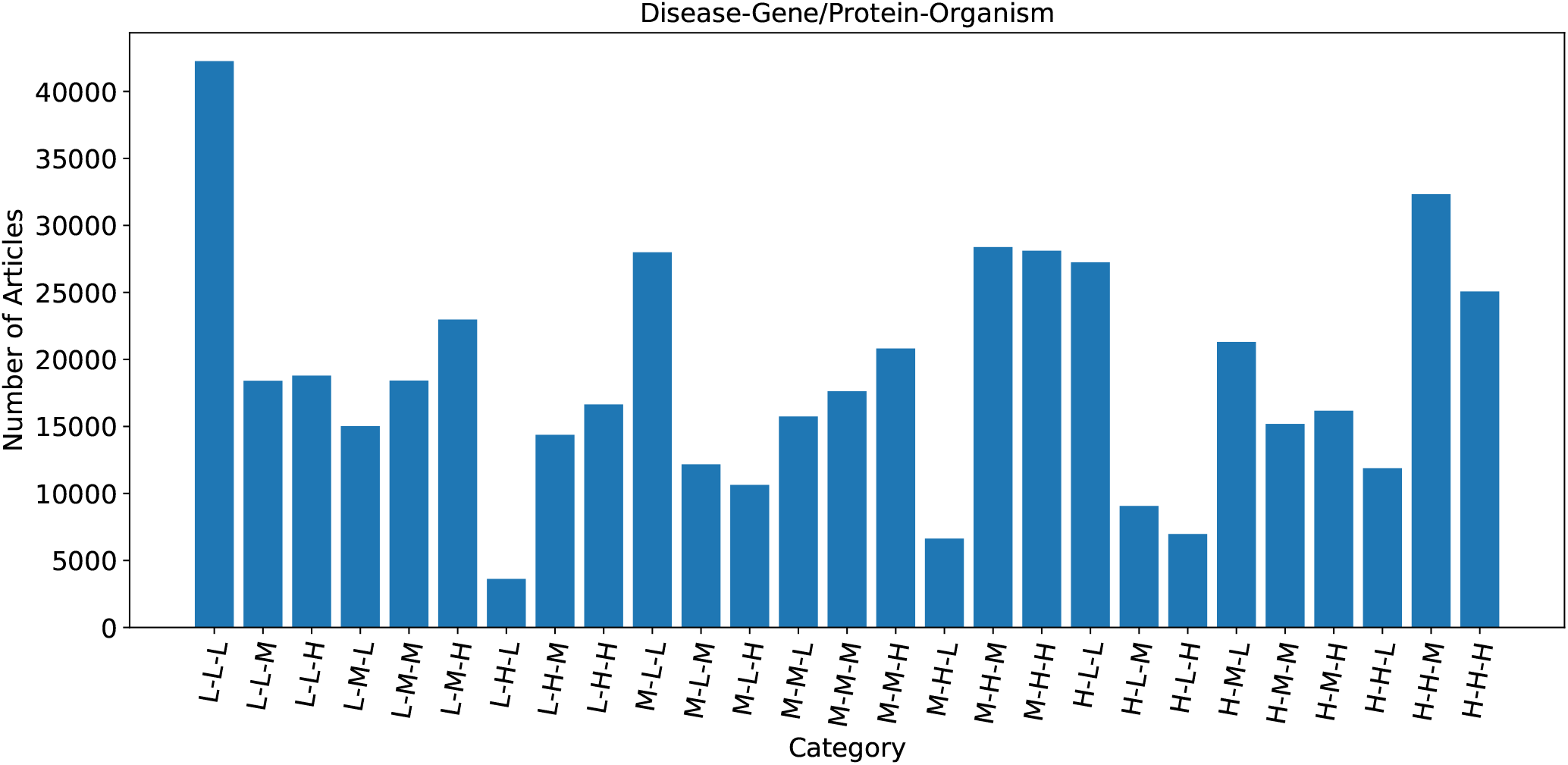
Distribution of articles based on the entity frequency. Here L, M, and H represent low frequency, medium frequency, and high-frequency tertile. The order of the label is Disease, Gene/Protein and Organism. For example, H-L-H represents articles that are high frequency for Disease and Organism and low frequency for Gene/Protein.

**Figure 5.**
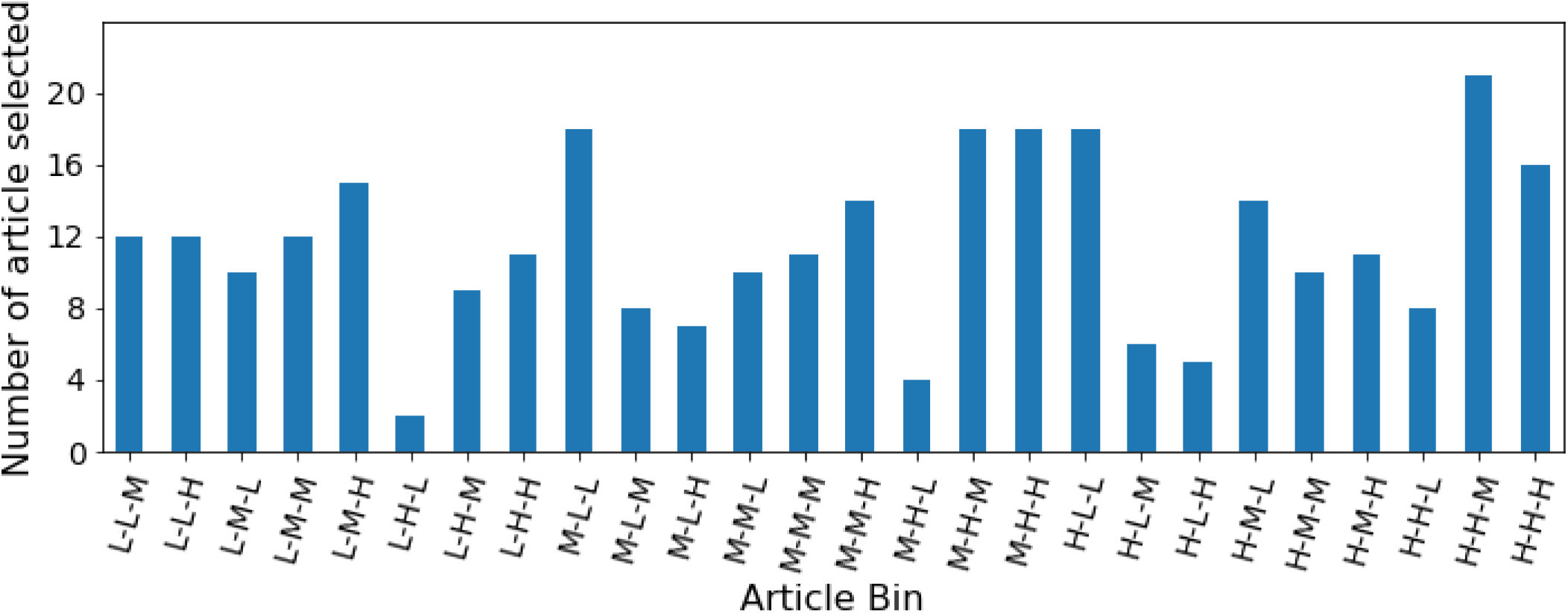
Number of articles selected from each bin for inclusion in the gold standard corpus of 300 articles. L, M, and H represent low frequency, medium frequency, and high-frequency tertile.

#### Ontology/terminology selection

The Europe PMC annotation pipeline currently uses a dictionary-based approach to tag Gene/Proteins, Diseases, and Organisms^1^. The term dictionaries are created from UniProt^5^, UMLS^30^, and the NCBI taxonomy^31^ for the Gene/Proteins, Diseases and Organisms, respectively. The pipeline annotates articles using predefined patterns and regular expressions to accommodate term variations from the dictionaries.

#### Gene/Protein

The Gene/Proteins dictionary is periodically generated from the SwissProt^32^ knowledgebase from the 2014 release. SwissProt is a manually reviewed resource of proteins and genes, and the knowledgebase is released in multiple formats. The entries in the Uniprot knowledgebase are structured to make it both human and machine-readable (for more details please follow https://www.uniprot.org/docs/userman.htm#convent). For tagging Gene/Proteins in the Europe PMC annotations workflow, the DAT file of the knowledgebase release is parsed, generating a Gene/Proteins dictionary from the gene name lines and their aliases (the gene name lines are denoted by starting the line with GN tag according to the knowledgebase data structure). The UniProt knowledgebase release, dated 2014, was used to generate the Gene/Proteins dictionary. In addition, a list of common English words (we call it a common-stop list) is used to avoid predominantly false-positive identifications, for example, ‘CAN’ as a gene name.

#### Disease

UMLS Diseases terms are used to create the Diseases dictionary. In UMLS, there are twelve different diseases/disorders (DISO) groups; four generate the Diseases dictionary because the other groups mainly comprise phenotypes and symptoms. The four DISO groups used are Disease or Syndrome (T047), Mental or Behavioural Dysfunction (T048), Neoplastic Process (T191), and Pathologic Function (T046). The ULMS version, dated 2015, was used to generate the Diseases dictionary.

#### Organism

The Organisms dictionary is based on the NCBI Taxonomy. Specific fields, such as acronym, BLAST name, GenBank common name and GenBank synonym, are used to populate the dictionary. The NCBI taxonomy version dated 2015 was used to generate the Organisms dictionary.

### Creation of annotation guidelines

A detailed concept annotation guideline is essential to develop a good corpus and resolve annotation disputes (Supplementary data file: annotation guideline). The CRAFT corpus provides comprehensive annotation guidelines^33^ explaining the annotation text spans and the entity type assignment. We based our annotation guidelines on the CRAFT corpus guidelines and expanded them to meet our requirements. A list of examples was included in the guidelines to assist curators. Before the start of the annotation work, a pilot study was conducted to annotate three articles. The aims of the pilot study are fourfold:

1. The pilot study helped curators estimate the workload to set timelines for the project;
2. Initial feedback was used to improve the annotation guidelines;
3. The curators familiarised themselves with the task and annotation tools;
4. It established the communication channel to manage the project.

### Article annotation

We worked with Molecular Connections ^1^, India, to employ three PhD level domain experts to annotate the corpus. We used a triple-anonymous approach to annotation; three annotators annotated the same articles independently to ensure annotation quality and validate inter annotation agreement. Annotation discrepancies were resolved by the majority vote to achieve/ensure the best quality annotation. That is, at least two annotators must agree on the annotation boundary and the entity type of the entity terms to pass the acceptance threshold. This maximised the total number of annotations. For example, if one annotator misses a term, it will likely be picked by the two other annotators. The triple-anonymous method made it possible to conveniently assess the inter-annotator agreements to ensure the annotation quality.

We sent the articles to the annotators in four batches. Between each batch, annotation quality and inter-annotator agreement were evaluated, and any confusion or quality issues were addressed. If necessary, updates to the annotation guidelines were made after each batch. To assess the quality of the annotations, the first batch consisted of only 30 articles, after which the number of articles per batch increased. This approach allowed us to resolve annotation discrepancies along the way and refine the annotator guidelines. Table 2 shows a detailed breakdown of these batches.

**Table 2.** Batch-wise annotation breakdown of articles and annotations.

|  | Annotator1 | Annotator2 | Annotator3 | Total | PMC count |
| --- | --- | --- | --- | --- | --- |
| Batch1 | 1583 | 1587 | 1587 | 4757 | 30 |
| Batch2 | 4745 | 4727 | 4733 | 14205 | 70 |
| Batch3 | 5604 | 5610 | 5611 | 16825 | 80 |
| Batch4 | 11932 | 11924 | 11931 | 36787 | 180 |

Annotators were instructed to view the articles on the Europe PMC website, where the existing dictionary-based annotations from Europe PMC text-mining pipeline are displayed using Scilite. The Hypothes.is annotation tool works as a layer on top of the Europe PMC website, allowing the curators/annotators to visualise and curate existing annotations and newly identified entity terms (Figure 6). We used Hypothes.is platform for annotations over other platforms such as BRAT^34^ and GATE^35^ as they require pre-processing of articles, for example, converting them to text files. Moreover, Hypothes.is provided easy access to Europe PMC website. We developed a set of standard schemes of tags for the curators to use and therefore classify the existing SciLite annotations.

**Figure 6.**
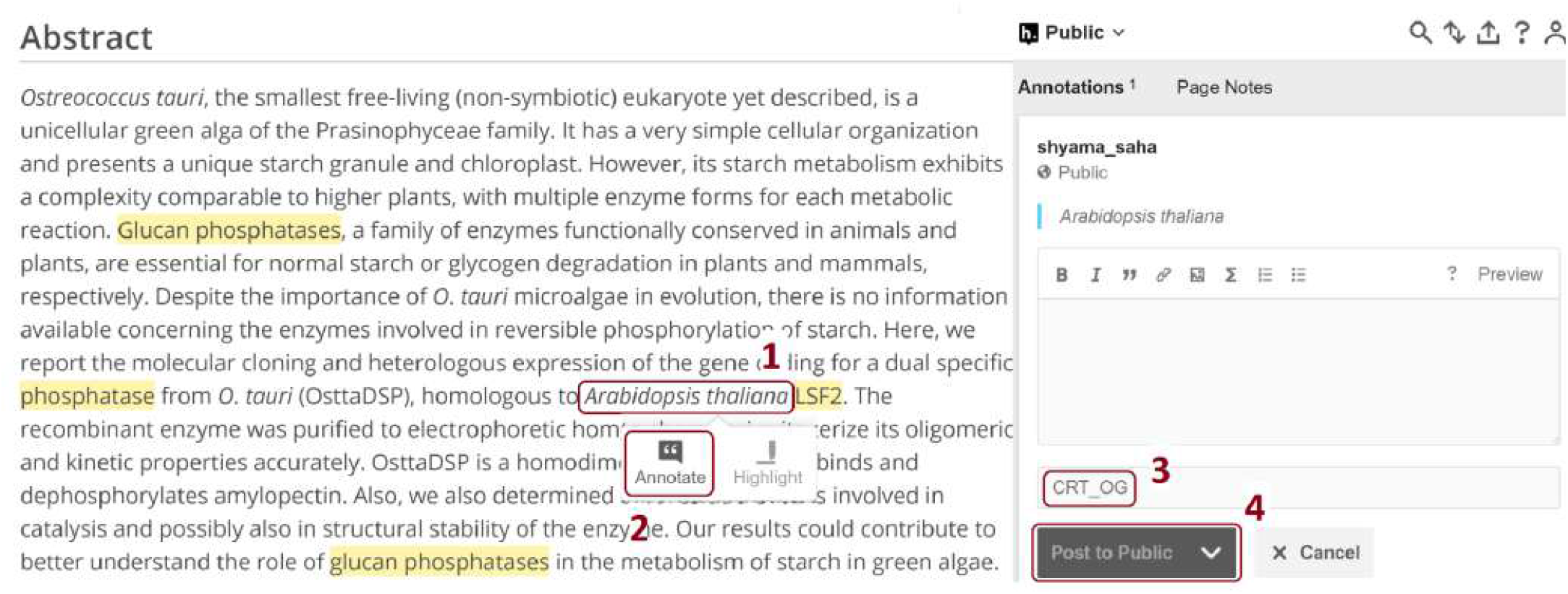
A screenshot of the Hypothes.is annotation platform overlayed on top of the Europe PMC website. Highlighted in yellow are existing dictionary-based text-mined terms. After selecting a term (1), users need to click the ‘Annotate’ button (2) to annotate the term. It will pop up the Hypothes.is annotation window on the right-hand side, allowing the annotators to add the annotation (3) and then save it using the ‘Post to Public’ button (4). Please refer to the supplementary material (Section ‘How to Use the Interface’ in the supplementary material “demo to molecular connections”) and Hypothes.is website for a detailed user manual.

The standard terms/tags were used as follows (Figure 7 shows an example of the use of these tags):

**Figure 7.**
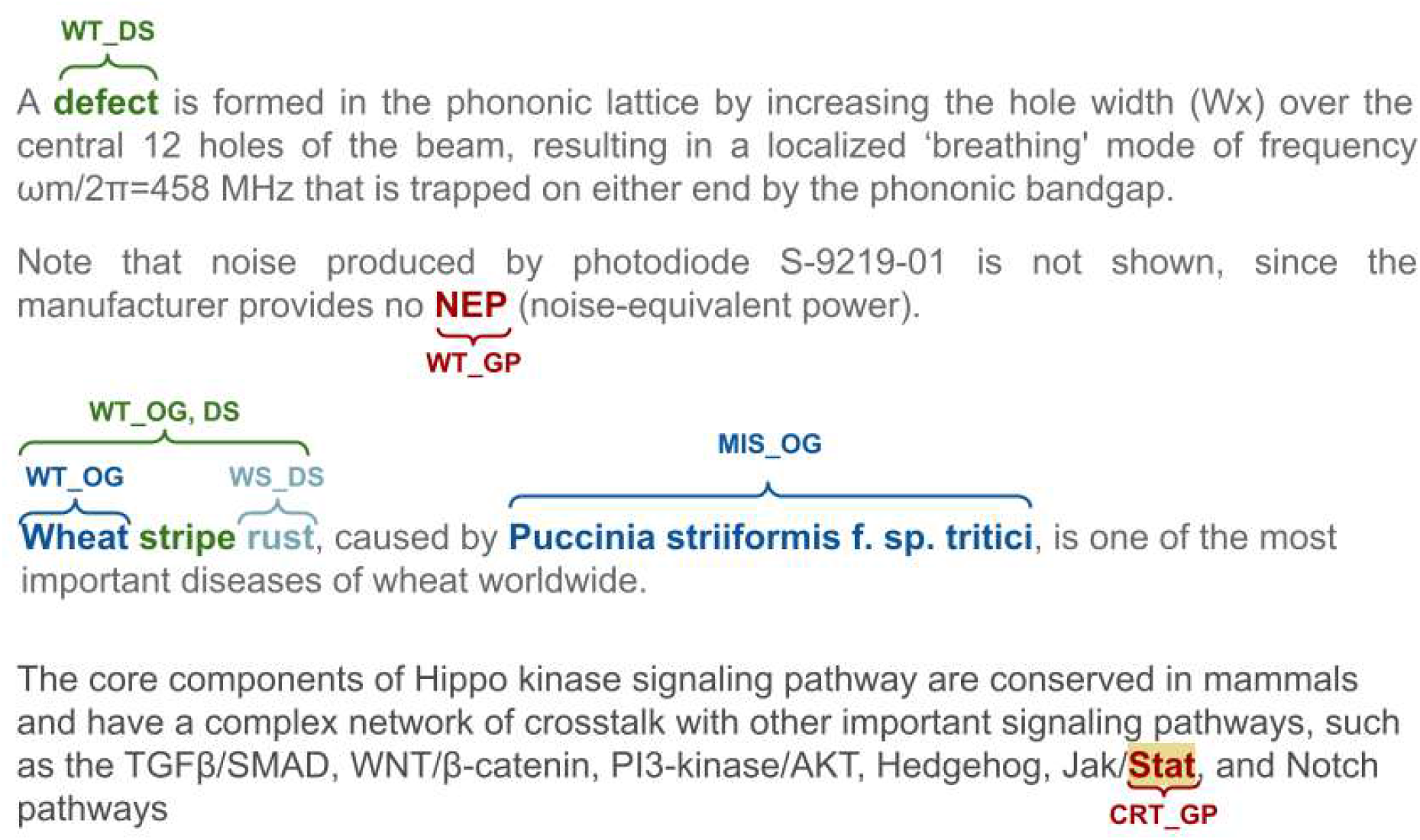
Example of human annotation correcting dictionary-based Europe PMC annotation using the tag set defined for this annotation task.

1. Correctness of annotation. Allows the annotators to verify existing Europe PMC annotations as Wrong Type (WT), Wrong Span (WS), Missing (MIS), or Correct (CRT).
2. Entity type. Three symbols were used to represent the entity types, GP for Gene/Proteins, DS for Diseases, and OG for Organisms.
3. A special tag ‘ALL’ allowed the annotators to apply the annotation of the current term to all occurrences of it across the article. This was useful in the case of reducing workload for the annotators and annotation cost but required additional work to find all the occurrences of a concept with an “ALL” tag in the post-processing phase.

These tags were used in combination to fully curate the annotations generated by the existing Europe PMC pipeline. For example,

- A correctly annotated Gene/Proteins (both entity type and annotation boundary) would be marked CRT_GP.
- A wrong Diseases annotation would be marked WT_DS; and if it had been for an organism; that would be marked as: [WT_DS][OG].

Although entity annotation was the main goal of this initiative, the annotators were asked to correct previously annotated Gene/Protein and Disease associations. Therefore, the third tag scheme for the association annotation, YGD, NGD, and AMB, represents a correct relationship, a wrong relationship and an ambiguous one, respectively.

A full list of all possible tags is in section 1.1 of the supplementary material: annotation guideline (Tag schema for annotation). Figure 8 shows the use of these tag sets for assessing the annotation discrepancies among three annotators.

**Figure 8.**
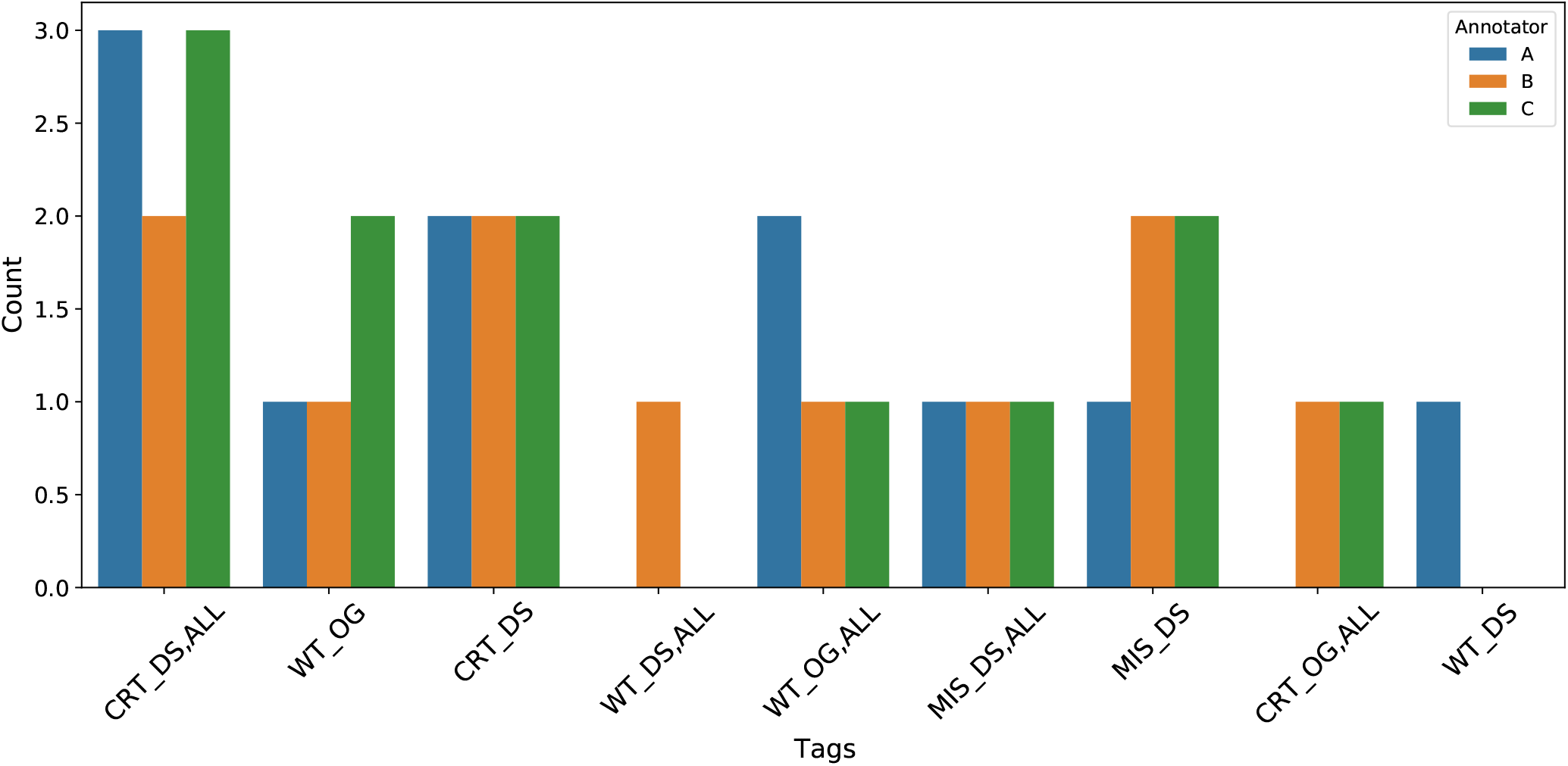
An example of the tag distributions from batch 1 showing the discrepancies between the annotators. Annotators used the ‘ALL’ tag to mark all mentions of the entity as correct (CRT) or wrong type (WT), missing (MIS), and so on. The DS and OG represent the Diseases and the Organisms entities respectively.

### Annotation extraction and processing

Hypothes.is^2^ is a free, open and user-friendly platform enabling annotation of web content. The annotators used Hypothes.is to highlight the span of the entity terms, add notes, and tag them with one of the available tags. They reviewed and marked pre-annotated terms as correct or incorrect and saved them using the Hypothes.is platform.

At Europe PMC, sentence boundaries are added to the article XML files using an in-house sentenciser prior to entity recognition. The Europe PMC text-mining pipeline annotates the bio-entities using a dictionary-based approach and displays them on the front-end HTML version via the web application (SciLite, which requires further processing of the annotated XML file). The Hypothes.is platform works on the front-end HTML version of the article. Each annotator set up a Hypothes.is account and thus their annotations were saved to the Hypothes.is server (Please refer to Section ‘How to Use the Interface’ in the supplementary material “demo to molecular connections” for detailed instructions). We retrieved the annotations using the Hypothe.is API in JSON format and it was converted to a CSV format using in-house tools. The Hypothes.is JSON reported the annotated terms and their locations with respect to the HTML version of the article.

The annotations from the JSON file were extracted or tagged in the sentencised XML file using regular expressions. However, due to the inconsistency between the HTML article page and the XML file, a small number of annotations could not be successfully extracted using regular expressions. We have identified that failure often occurs when an annotation is in a table. We post-processed the Hypothe.is JSON files for presenting the corpus to the wider community in multiple formats. More details are in the following sections. Figure 9 shows an overview of the process.

**Figure 9.**
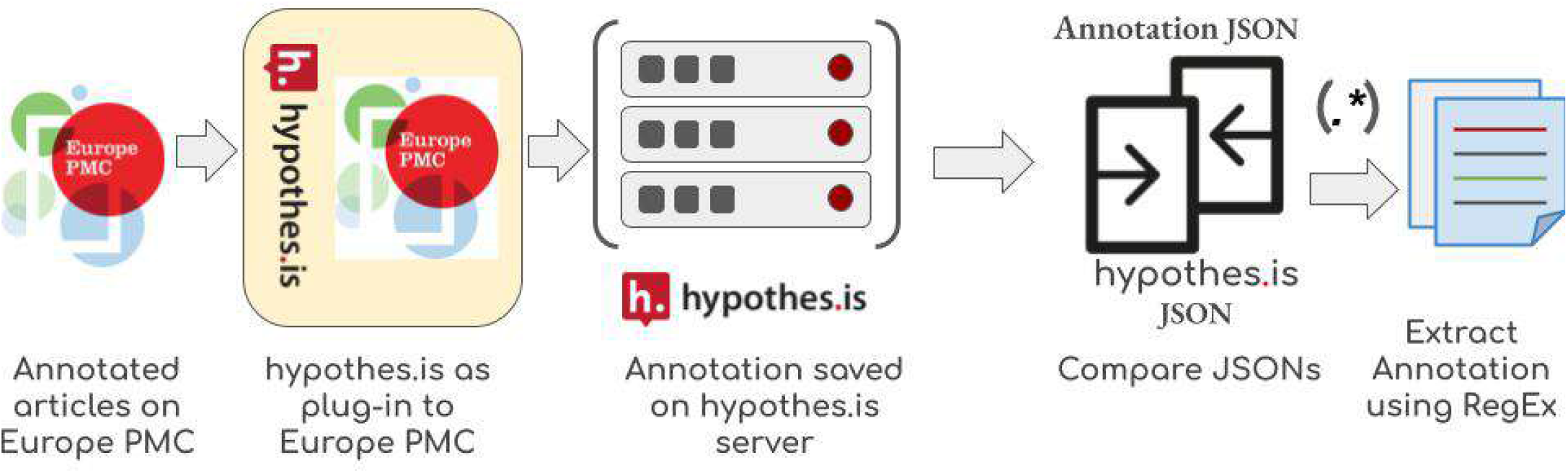
Annotation extraction workflow. Hypothes.is was added onto Europe PMC as a plug-in for the annotation work. Annotators saved their annotations to the Hypothes.is server in JSON format and it was retrieved and converted to CSV format using in-house tools. Europe PMC parses the XML version of the articles for sentence tagging and annotating named entities and displays an HTML version on the front end. We compared the hypothe.is annotation JSON files against the XML version and extracted the annotations using regular expressions.

### Data Records

We have made this training set available in the following formats at:

https://gitlab.ebi.ac.uk/literature-services/public-projects/europepmc-corpus

1. Stand-alone curator annotations.
  a. CSV
  b. JSON
  c. Inside-outside-beginning (IOB)
2. Full-text XML files (without EPMC annotations)
3. Full-text XMLs with sentence boundary (we add <SENT> tag to annotate the sentence boundary)
4. Europe PMC annotation in JSON format.

To fit the diverse needs of the annotation users, the corpus provides multiple formats of annotations from the raw annotations of Hypothes.is platform (in CSV format) to the standard and ready to use IOB format. In addition to the annotations, original full-text articles are released in XML format without the tags. With the raw annotations in CSV format and full-text XML files, researchers can apply their own text-mining tools to extract the annotations. The comma-separated values (CSV) raw annotation files contain three fields (exact, prefix, and suffix) that are critical to locating the human annotations. “exact” is the annotation itself while “prefix” and “suffix” are characters before and after the annotation, respectively. By combining “prefix”, “exact”, and “suffix”, the snippet can locate the annotation using regular expressions. Raw annotations from all three human annotators are available on GitLab, which are helpful for studies of agreement between annotators. Annotations in JavaScript Object Notation (JSON) and IOB formats are provided in addition to raw annotations. Both JSON and IOB format annotations are preprocessed so that only annotations agreed on by at least two annotators are included. The IOB format provides sentences with IOB tags and follows the CoNLL NER corpus standards^36^. While the IOB format is widely used in named entity recognition (NER), researchers may prefer other tagging formats so the JSON format provides sentences and annotations for researchers that are interested in transforming annotations into other tagging formats. Full-text articles are also available in the format that articles are split into sentences by the Europe PMC text mining pipeline.

### Technical Validation

This paper presents a corpus of 300 full-text open access articles from the biomedical domain, human-curated with the entities Gene/Proteins, Diseases, and Organisms. Eight articles from the corpus do not contain any entity annotations because the human annotators removed existing dictionary-based annotations as false positives. These articles came from 5 different bins. Table 3 and 4 show an overview of the human-annotated terms and compares these to the existing Europe PMC dictionary-based approach. To evaluate the dictionary-based approach, we applied majority voting acceptance criteria on the granular level annotation tags, that is, entity type tags (GP, DS, OG) along with the correctness tags (CRT, MIS, WT, WS).

**Table 3.** Overall annotation statistics comparing the existing Europe PMC dictionary-based text mining approach to the human curation for the selected 300 gold standard articles. Overall we have gained around 11k term annotations, with the highest gain existing for the Gene/Protein category. We report unique term count based on the string match and how many normalise to a database identifier of the databases mentioned above rather than unique database identifier counts.

|  |  | Europe PMC dictionary-based |  |  |  | Gold standard human annotation |  |  |  |
| --- | --- | --- | --- | --- | --- | --- | --- | --- | --- |
|  |  | Gene/<br>Protein | Disease | Organism | Total | Gene<br>/Protein | Disease | Organism | Total |
| Annotations | Total | 28,869 | 10,515 | 18,040 | 57,425 | 36,369 | 14,518 | 21,491 | 72,378 |
|  | Unique | 3,419 | 1,752 | 1,700 | 6,871 | 5,600 | 2,037 | 2,347 | 9,970 |
| Normalised<br>to a DB entry | Total | - | - | - | - | 21,664 | 8,476 | 16,021 | 46,161 |
| Median per<br>article | Total | 53.5 | 19.5 | 34 | 170 | 54.5 | 16 | 30 | 192 |
|  | Unique | 12 | 8 | 8 | 36 | 13 | 6.5 | 8 | 44.5 |
| Max annotation<br>per article | Total | 722 | 219 | 407 | 955 | 795 | 478 | 456 | 940 |
|  | Unique | 113 | 78 | 111 | 156 | 178 | 76 | 170 | 201 |

**Table 4.**
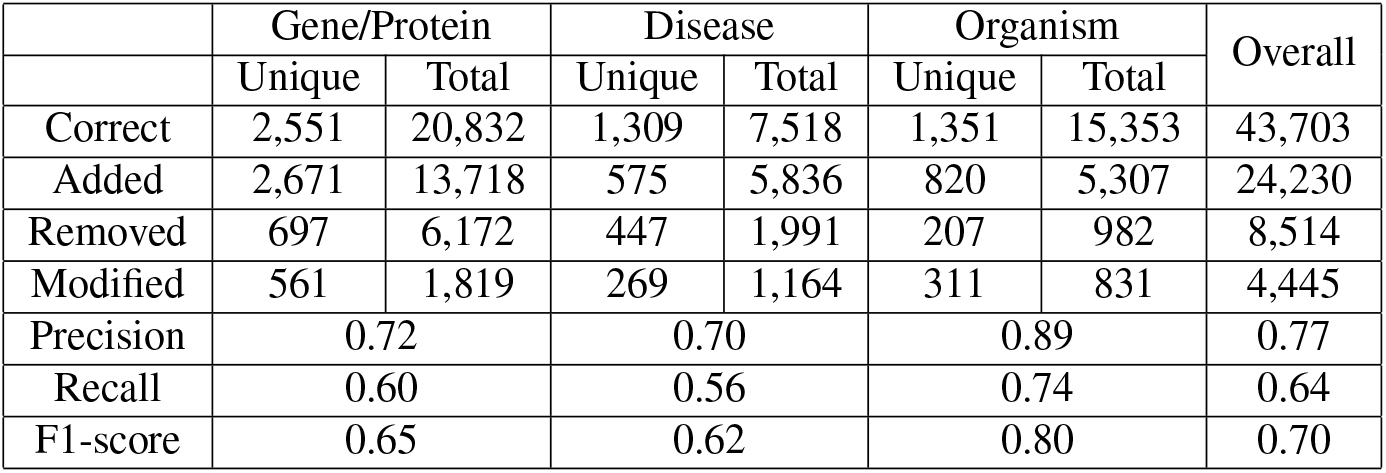
Evaluation of current Europe PMC dictionary workflow against the human annotation. This table shows the number of dictionary-based Europe PMC annotations updated by human annotators. A large proportion of the Europe PMC annotations are confirmed as correct by the human annotators, although they also added/annotated a significant number of previously unidentified/unannotated terms. The Europe PMC pipeline misses a proportion of these terms due to outdated dictionaries. The removed terms are often common English words or short acronyms. Gene/Protein terms (GP) are more likely to be removed than other entity types, that is, Diseases (DS) and Organisms (OG), due to the frequency of occurrence and the false positive rate for three-letter gene-protein acronyms. This row also counts the annotation where the dictionary-based approach wrongly assigned the type (WT), e.g. Diseases entities wrongly tagged as Gene/Proteins (WT_GP) by the Europe PMC dictionary-based approach (annotators used WT_GP, DS tag) will be added to the ‘removed’ cell count for the Gene/Proteins and ‘added’ cell for the Diseases. The “Modified” row shows the number of entities that were modified/split into multiple entities (WS). The overall column is the summation of correctness tags (CRT), i.e. CRT, Missing (MIS) and Wrong Span (WS), going under the Correct, Added and Modified rows. For the WT tag, they were split into two, one under the Removed column and the rest under the Modified row. When an annotation is assigned WT_GP, it means that it is a wrong Gene/Proteins annotation and removed from the annotation set, whereas the [WT_GP, DS] tag means the annotation was not removed from the annotation set, but the entity type is modified.

The triple-anonymous annotation approach had an overall inter-annotator agreement of 0.99. At this level, we assigned granular tags to appropriate entity types. For example, CRT_GP and WS_GP tags were mapped to the GP tag and used the strict evaluation rule defined by the Message Understanding Conference (MUC) for the inter-annotator agreement. High inter-annotator agreement with the strictest methods shows that most of the annotations were agreed upon by all three annotators (Table 5). A total of 767 annotations were discarded because just one annotator annotated them. Among these discarded annotations, 289 annotations had overlapping text spans, with the 1,005 annotations agreed upon by two annotators. For example, two annotators annotated “Welsh Mountain sheep”. However, the third annotator only annotated “sheep” from “Welsh Mountain sheep”. Both of them are correct in terms of the definition of species. Only 478 annotations were truly discarded, accounting for 0.7% of total annotations. Further inspection of the discarded annotations may validate some and help keep the correct ones, but we did not consider this to be a major blocking task.

**Table 5.** Inter-annotator agreement statistics. We evaluated annotation agreement using SemEval-2013 Task9.1 strict rule. According to the strict evaluation rule, an annotation agreement is reached only when two annotators agree on the term span and the annotation type. We achieved an overall agreement of 0.99. The first row of this table shows the entity-level breakdown of annotations that were rejected due to the voting system, i.e. at least two annotators must agree on the annotation term, boundary and entity type. Some of these entities were annotated by the other annotators with different entity boundaries.

| Agreed by | Gene/Protein | Disease | Organism | Overall |
| --- | --- | --- | --- | --- |
| 1 annotator | 270 | 178 | 319 | 767 |
| 2 annotators | 480 | 309 | 216 | 1005 |
| 3 annotators | 35934 | 14237 | 21298 | 71469 |

Our analysis of the distribution of tags set (Figure 10) shows the highest number of missing terms by the dictionary-based approach is from the Gene/Proteins type (MIS_GP tag). This might be due to the fact that our Gene/Proteins dictionary was last updated in 2014. Updating an entity dictionary involves a number of manual human edits, making it difficult to maintain. Although we were aware of the limitations of the common-stop list approach to limiting false positives, human annotation showed only a small number of these terms (1.6% tagged as MIS_GP) were inappropriately excluded. Using this gold standard data to train the state-of-the-art machine learning/deep learning models for entity recognition eliminates these challenges. We observe the same trend for the false-positive identifications, i.e. WT_[GP|DS|OG]. The highest number of false positives are from the Gene/Proteins type followed by the Diseases and Organisms terms, respectively. The wrong-type annotation counts are quite low; annotators only correct the entity type for a small number of annotations. This perhaps reflects the way the Europe PMC annotation pipeline works. This pipeline applies dictionaries sequentially, first the Gene/Proteins dictionary, followed by the Diseases dictionary, and then the Organisms dictionary. Once an entity is tagged, it becomes unavailable to tag with subsequent dictionaries, likely reducing false-positive Diseases and Organisms entity identifications. Our analysis shows only a few terms were assigned to the wrong entity type due to this approach, proving our sequential method works. Table 6 shows how many term annotations were updated to reassign the entity type.

**Table 6.** Europe PMC dictionary-based entity annotation follows a sequential manner to annotate the entities. For example, we apply the Gene/Proteins dictionary before the Disease dictionary, making the Gene/Protein terms unavailable for the disease tagger. We minimise the false positive identifications through this approach. This table shows the number of wrong entity type assignments by the Europe PMC approach corrected by the manual annotators. Europe PMC misses a small percentage of the Disease and Organism entities due to the sequential approach. We are showing Europe PMC annotation in the rows and the manually corrected ones in the columns.

|  | human annotation |  |  |  |
| --- | --- | --- | --- | --- |
| Europe PMC Annotation |  | Gene/Protein | Disease | Organism |
|  | Gene/Protein | - | 324 | 113 |
|  | Disease | 47 | - | 18 |
|  | Organism | 19 | 110 | - |

**Figure 10.**
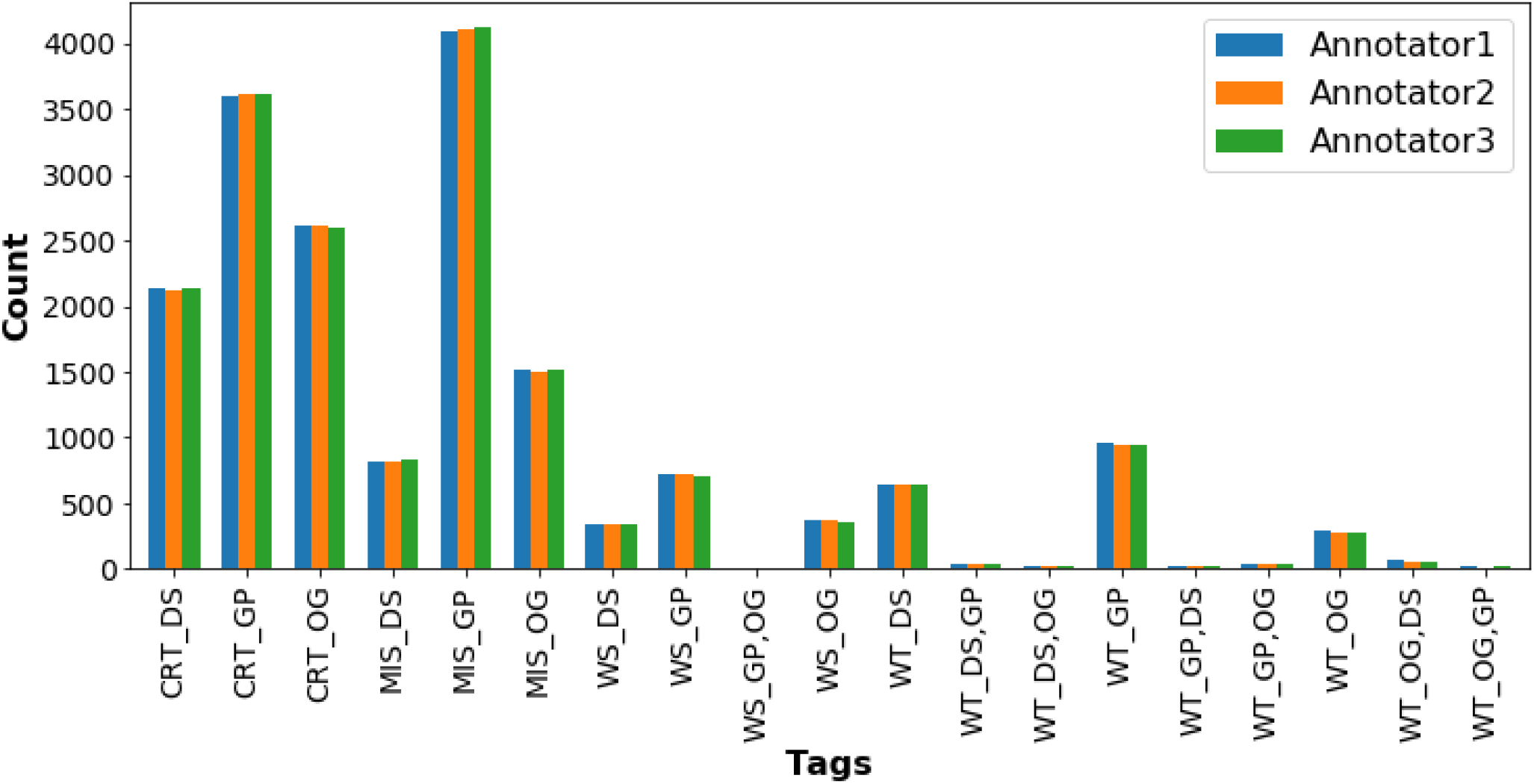
Entity tags distribution of the corpus and the comparison among the annotators. A large number of Gene/Proteins terms are missed by the dictionary annotation. This figure demonstrates high inter-annotator agreement; correct (CRT), missed (MIS), wrong span (WS), and wrong type (WT). The latter part of the tag represents the entity type namely, Disease (DS), Gene/Protein (GP), and Organisms (OG). Annotators use the WT keyword to remove an annotation and to change the entity type of annotation. They submit the correct entity type by adding the correct entity type keyword after the WT tag, e.g. WT_OG, DS.

The special ‘ALL’ tag was used to indicate that the annotation of a term applies to all occurrences of the term within the article. This was a significant time-saver for articles that mention a particular entity tens or 100s of times. A total of 23,281 (7,336 unique) terms were tagged ‘ALL’.

Because Hypothes.is allows free text in the tag field, we identified a small number of errors in the tag names; for example, ten annotations from annotators 1 and 2 use ‘DIS’ instead of ‘DS’; one annotation uses ‘CRt’ instead of ‘CRT’. We corrected these errors for downstream analysis.

The titles of sections within a research article can vary widely but typically fall into a small number of categories. For example, “Methods” and “Methods and Reagents” are both classed as Methods sections. In the Europe PMC annotation pipeline, section titles are normalised to a set of 17 titles^37^. Figure 11 shows the entity distribution across these sections. As anticipated, we found a high frequency of entity mentions in an article’s main sections, which demonstrates the value of full-text annotation versus using only abstracts^38^. This entity distribution may help design a targeted annotation approach when resources are limited.

**Figure 11.**
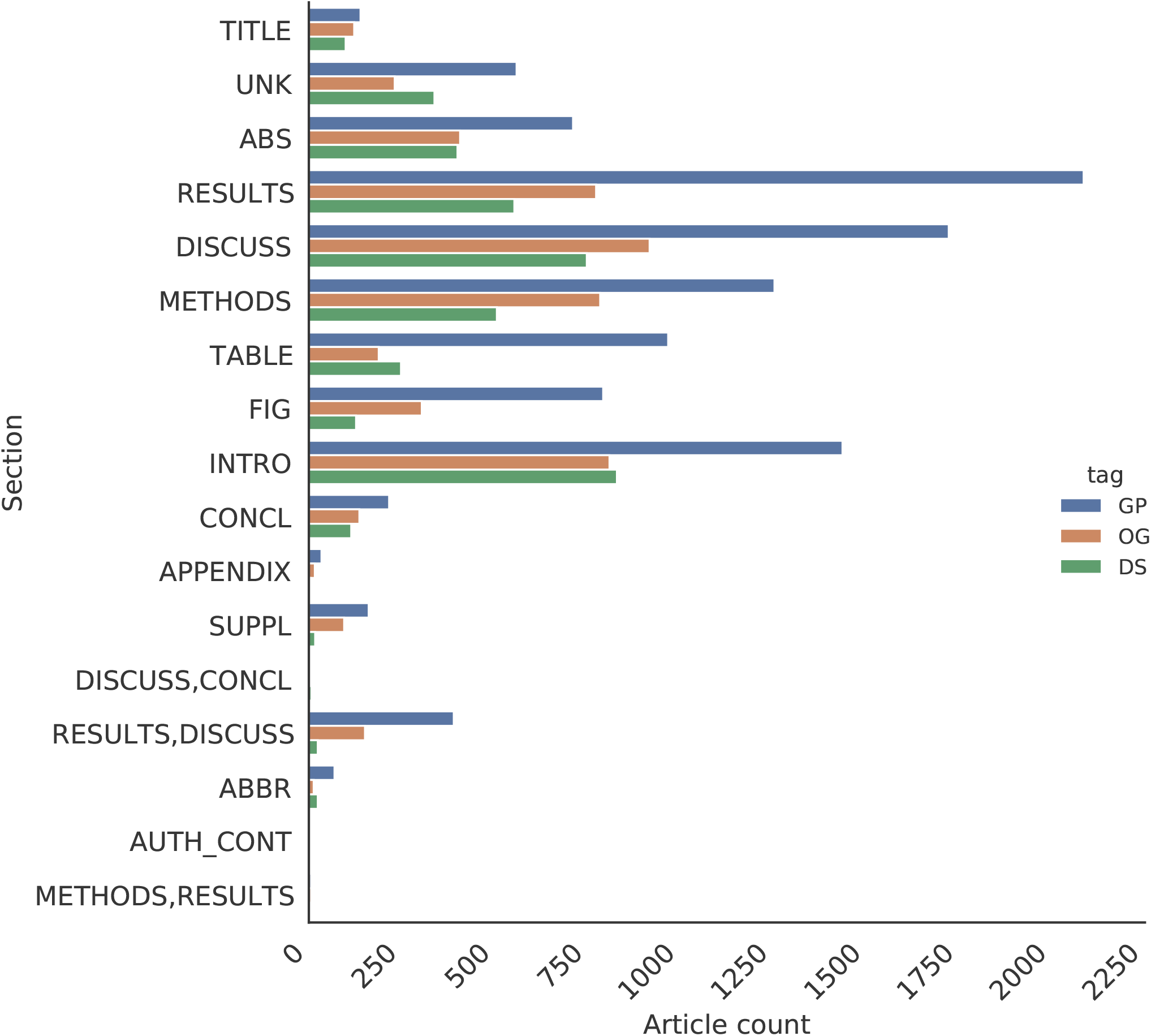
Term frequency distribution across different sections. The result, discussion, method, and introduction sections contain the highest number of entity mentions. Gene/Proteins mentions in tables and figure titles are significantly higher than Diseases and Organisms mentions.

## Discussion and Conclusion

Gold standard datasets are essential for capitalise on the revolutionary deep learning methods for NLP and text-mining. Although a number of gold standard datasets are available for biomedical entities, all except the CRAFT corpus and NLM-Chem are based on abstracts. As demonstrated here, high numbers of the entities appear in the body of the articles. Moreover, the abstract is often a summary of the rest of the article and has very different language constructs. These differences make the gold standard abstract datasets unsuitable for full-text article annotation tasks.

The EPMCA corpus is larger than the CRAFT corpus and annotates Gene/Protein, Disease, and Organism entity types and comprises ≈ 1000 sentences with Gene/Protein and Disease association annotations, making it a unique addition to this domain. The approach to construction is also slightly different. We used a triple-anonymous approach rather than the lead annotator approach used for the CRAFT corpus and we achieved a very high inter-annotator agreement across the corpus and all entity types. The use of the pre-annotated articles for curation is another important feature of this dataset, as it allowed us to start a large set of initial annotations to validate at the same time as the annotators identified new annotations. We hope that the two approaches complement each other to support the future development of novel deep learning approaches.

We are currently using this corpus to train deep learning methods to address the challenges mentioned above for named entity recognition (NER). Moreover, we have achieved a better sentence co-occurrence based target–disease association identification through the improved NER system. The association annotation process is being further improved using the association annotation dataset of the corpus. We hope this dataset will be just as beneficial to the community.

## Supporting information

demo to molecular connections

annotation guideline

## Code availability

The code is available at the repository https://gitlab.ebi.ac.uk/literature-services/public-projects/europepmc-corpus/. The scripts include cleaning and formatting the annotations from Hypothes.is platform and generates the dataset in IOB format for input to deep learning algorithms.

## Acknowledgements

We thank the curators at Molecular Connections (https://www.molecularconnections.com/) for their biocuration efforts. We would also extend our thanks to Heidi J. Imker from University of Illinois at Urbana-Champaign and Melissa Harrison Matt Jeffryes, and Mariia Levchenko from team Europe PMC for their valuable suggestions. Last but not least, we would like to thank Europe PMC and Open Targets for strategic guidance on the development of the Europe PMC corpus. This work was supported by: the European Molecular Biology Laboratory-European Bioinformatics Institute (X.Y, A.V); Europe PMC grant, provided by 37 funders of life science research (https://europepmc.org/Funders/) under Wellcome Trust Grant 221523 (A.V, V.V) and the OpenTargets grant 2056 (S.T, S.S).

## Author contributions statement

X.Y. designed and developed annotations guidelines, conceived the experiment(s), analysed the results and wrote the initial draft of the manuscript. S.S. contributed to writing the manuscript and analysing the dataset. A.V. contributed towards annotator guidelines, annotator tags and data structure in Hypothes.is. S.T. developed scripts for generating machine-trainable IOB formatted datasets, contributed to proofreading, writing the paper, analysed the data and regenerated all the figures for reproducibility. V.V collected and sampled data from Europe PMC database. J.M. conceived the idea and supervised the development of the dataset. All authors reviewed the manuscript.

## Footnotes

1 https://molecularconnections.com/

2 https://web.hypothes.is/

