## Supplementary material for "Europe PMC Annotated Full-text Corpus for Gene/Proteins, Diseases and Organisms": demo to molecular connections

### Annotate the text using unified names

“Span” indicates the span (i.e. the set of characters to select) of an entity. It refers to a selection of consecutive characters of the entity. Annotations in **Blue** denote disease and in **Green** denote organism.

1. Annotation is correct for both the span and type

   Finally, the researchers report that injection of PKHB1 reduced the **tumor** burden in a mouse model of CLL. [PMC4348493]

   “tumor” is annotated as Disease, which is correct both for span and type.

2. Annotation type is correct but the span is wrong

   Finally, the researchers report that injection of PKHB1 reduced the **tumor burden** in a mouse model of CLL. [PMC4348493]

   “tumor burden” is annotated as Disease. The correct annotation should be “tumor” and the type should be Disease. The annotation is longer than the expected entity “tumor”. Therefore, it has wrong span but correct type.

   Finally, the researchers report that injection of PKHB1 reduced the **tum**or burden in a mouse model of CLL. [PMC4348493]

   “tum” is annotated as Disease. The correct annotation should be “tumor” and the type should be Disease. The annotation is shorter than the expected entity “tumor”. Therefore, the annotation has wrong span but correct type.

3. The span is correct but the type is wrong

   Finally, the researchers report that injection of PKHB1 reduced the **tumor** burden in a mouse model of CLL. [PMC4348493]

   “Tumor” is annotated as Organism. The correct annotation should be “tumor” and the type should be Disease. Therefore, the annotation has wrong type but correct span.

4. Both the span and type are wrong

   Finally, the researchers report that injection of PKHB1 reduced the **tumor burden** in a mouse model of CLL. [PMC4348493]

   “tumor burden” is annotated as Organism. The correct annotation should be “tumor” and the type should be Disease. Therefore, the annotation has wrong type and wrong span.

5. Missing entity (false negative)

   Finally, the researchers report that injection of PKHB1 reduced the tumor burden in a mouse model of CLL. [PMC4348493]

   “tumor” is missing from the annotation. Therefore, it’s a missing annotation.

Gene-Disease Relationship annotation:

1. The relationship is correct
   a. Both of the gene and disease entities are correct
   b. The relationship exists between the entities.
2. The relationship is wrong
   a. One or both of the entities in the pre-annotated relationship have wrong type
      i. Refer to the entity annotation to check whether the type is correct
   b. Entities are correct but relationship doesn’t exist
3. The relationship is ambiguous:
   a. Both entities have correct type, but the relationship is ambiguous
4. In the current phase, it is not necessary to annotate missing relationship

### Tag scheme for annotations

1. Tags for indicating wrong/correct annotations:

| **Category** | **Tag** |
|---|---|
| Wrong type | WT |
| Wrong span | WS |
| Missing | MIS |
| Correct | CRT |

Table 1

2. Tags for entity:

| **Name** | **Tag** |
|---|---|
| Gene/Protein | GP |
| Organism | OG |
| Disease | DS |

Table 2

3. Tags for gene-disease relationship:

| **Category** | **Tag** |
|---|---|
| Correct relationship | YGD |
| Wrong relationsjip | NGD |
| Ambiguous | AMB |

Table 3

4. Special tag:

| **Special Tag** | **Tag** |
|---|---|
| All | ALL |

Table 4

### Usage of annotation tags

In order to indicate both the wrong correct tags. We suggest to use following scheme to report wrong/correct/missing annotations.

A. Annotation is correct for both the span and type

| **Type** | **Tag** |
|---|---|
| Gene/Protein | CRT_GP |
| Organism | CRT_OG |
| Disease | CRT_DS |

Table 5

B. Annotation type is correct but the span is wrong

| **Type** | **Tag** |
|---|---|
| Gene/Protein | WS_GP |
| Organism | WS_OG |
| Disease | WS_DS |

Table 6

C. The span is correct but the type is wrong

In order to record the wrong annotation type, we need to use underscore to indicate the wrong type. For example, [WT_GP] means the wrong annotation type is Gene/Protein. The correct type can be indicated using an additional tag as shown below. If the annotation is a false positive, then we don't need to provide the correct type.

| **Wrong Type** | **Correct Type** | **Tag** |
|---|---|---|
| Gene/Protein | Organism | [WT_GP][OG] |
| Gene/Protein | Disease | [WT_GP][DS] |
| Gene/Protein | None | [WT_GP] |
| Organism | Gene/Protein | [WT_OG][GP] |
| Organism | Disease | [WT_OG][DS] |
| Organism | None | [WT_OG] |
| Disease | Gene/Protein | [WT_DS][GP] |
| Disease | Organism | [WT_DS][OG] |
| Disease | None | [WT_DS] |

Table 7

D. Both the span and type are wrong

Refer to Table 7. If the type is wrong, the span is not important. Therefore, we don't need to record whether the span is right ot not. Use the scheme in Table 7.

E. Missing entity (false negative)

| **Type** | **Tag** |
|---|---|
| Gene/Protein | MIS_GP |
| Organism | MIS_OG |
| Disease | MIS_DS |

Table 8

F. Usage of the special tag

The special tag [ALL] is used when the current annotation can be applied to the same annotations in the full text. For example, if all the pre-annotations of “tumor” are correctly tagged as Disease with the right span in one article, then we can use the combination of [CRT_DS][ALL] to indicate all the same pre-annotations o “tumor” are correct. Therefore, we can skip the same pre-annotations after it.

G. Annotation of gene-disease relationship

If the pre-annotation of the relationship is correct, use tag **YGD** from Table 3.

If the pre-annotation of the relationship is wrong, use tag **NGD** from Table 3:

- One or both the entities have wrong type
- Both entities have correct type, but no relationship

If the pre-annotation of the relationship is vague/ambiguous, use tag **AMB** from Table 3:

- Both entities have correct type, but the relationship is ambiguous

#### How to use the interface

The following examples illustrate how to use the Hypothes.is plug-in. Chrome must be installed as the current plug-in only support Chrome. The screenshots may differ from the aforementioned tagging scheme, therefore please refer to tagging scheme for annotation.

1. Annotators create Hypothes.is account.
2. An invitation of joining the annotation group will be sent to all annotators.
3. Install Hypothes.is plug-in in Chrome app store (Add to Chrome)

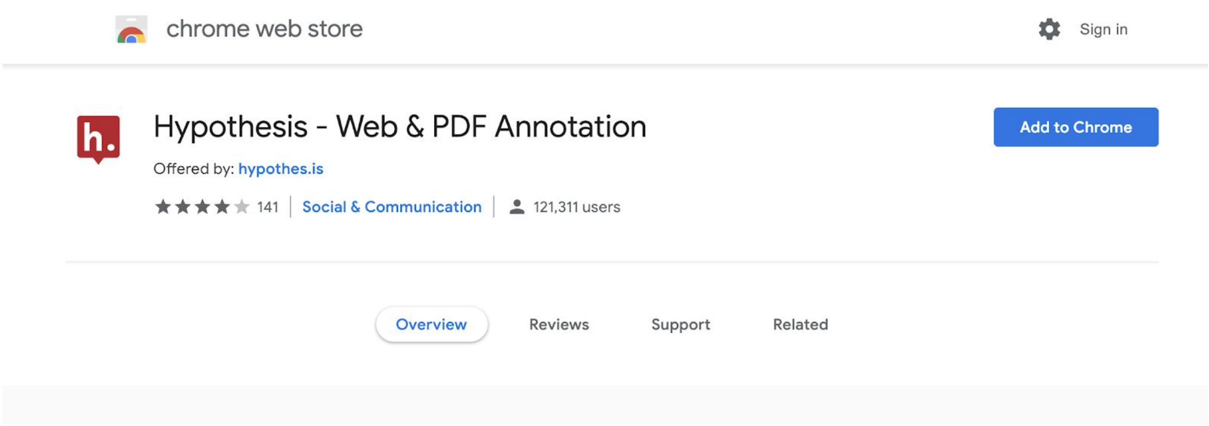

4. Open an article in EuropePMC using PMCID

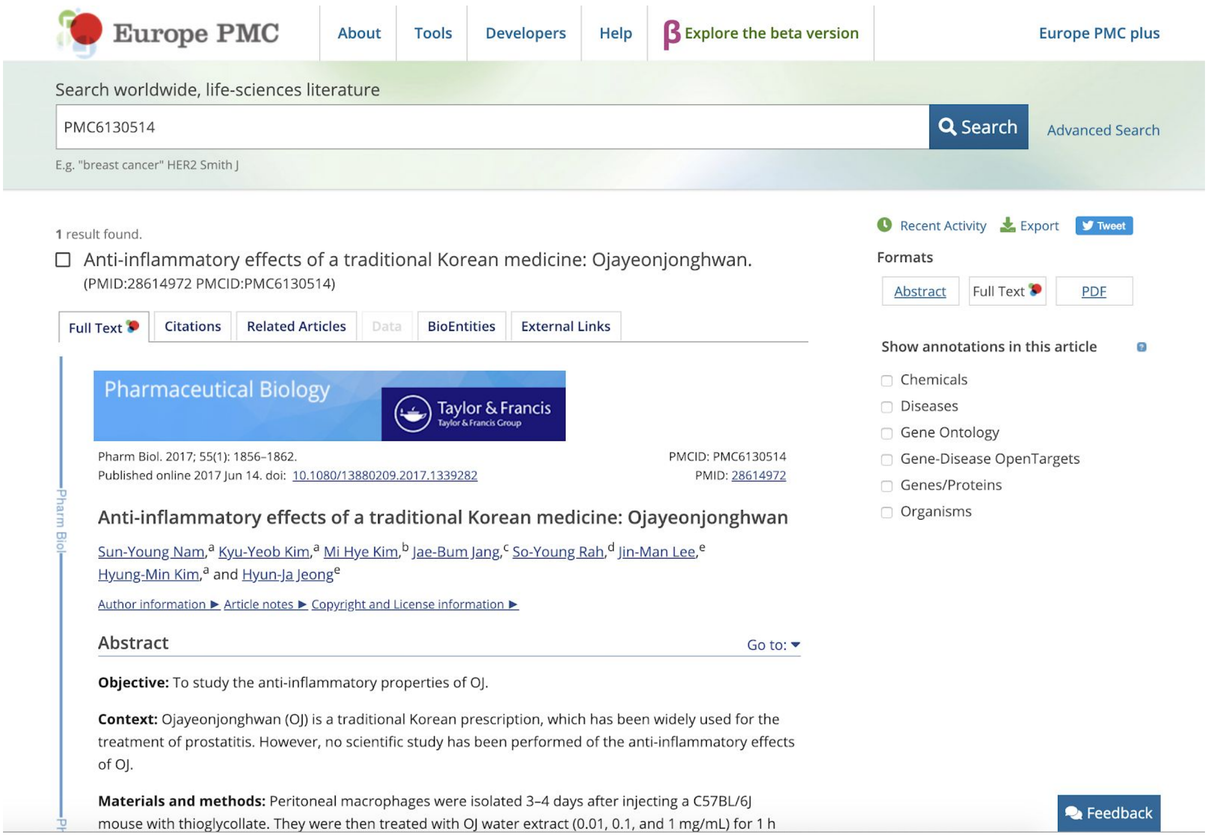

5. Select Gene/Protein, Disease, Organism and Gene-Disease OpenTargets ( if available) from the right panel to show annotations

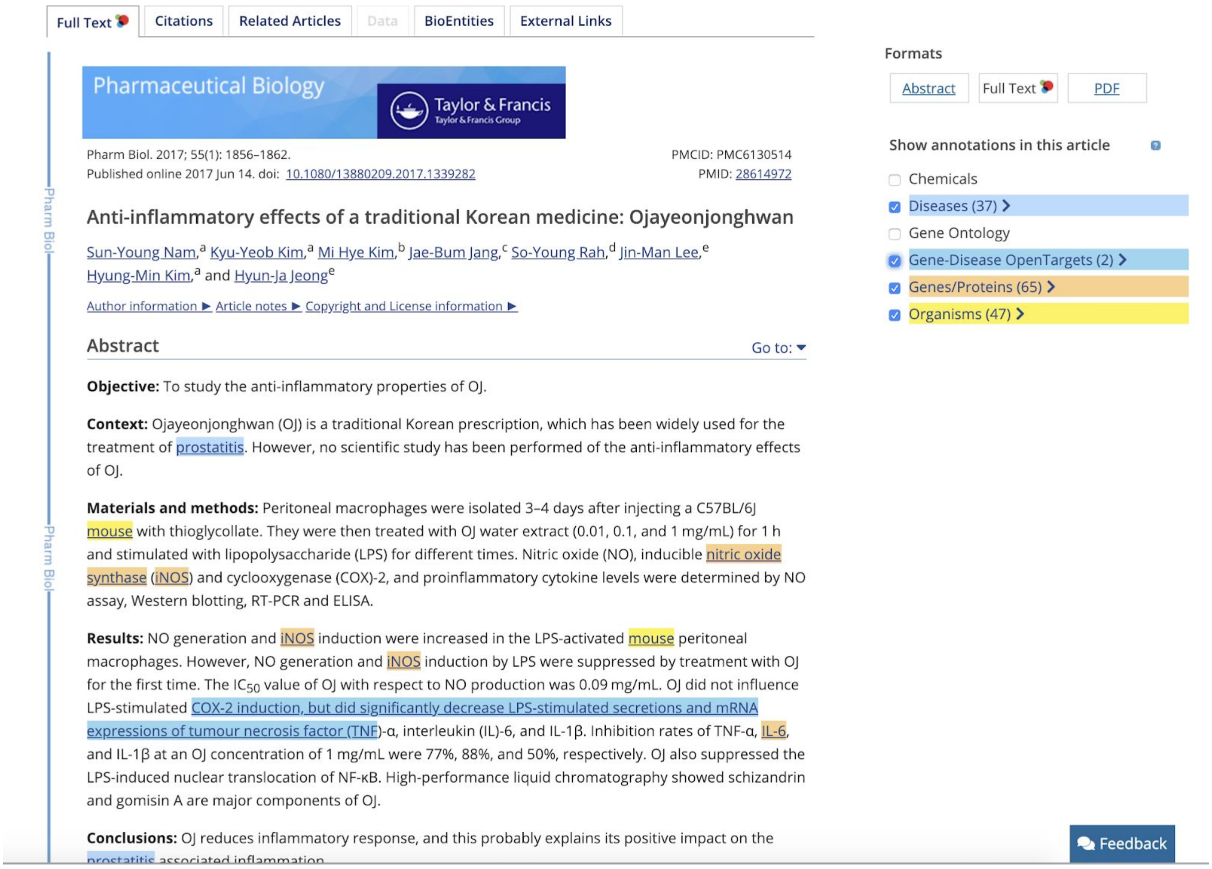

6. Click the Hypothes.is plug-in symbol to activate Hypothes.is

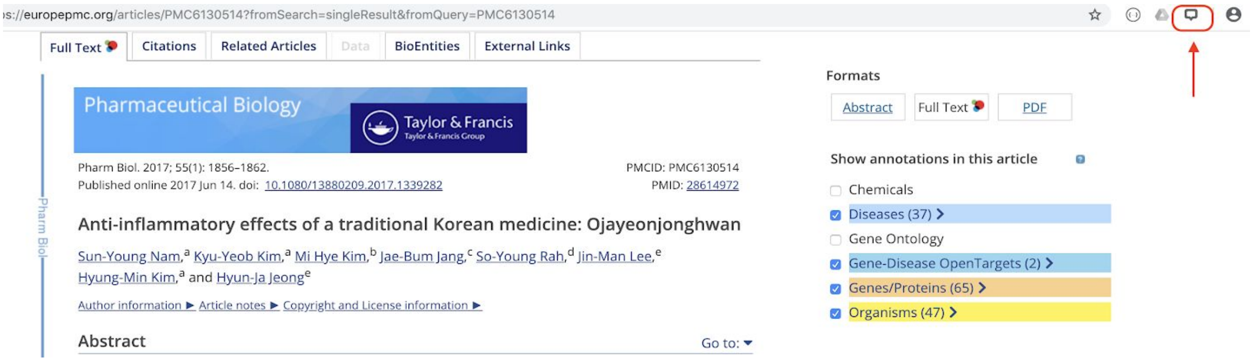

7. The pre-annotated text are highlighted in different colours. Click the highlighted text, a window will pop up to show more details e.g. the entity type and the annotated text.

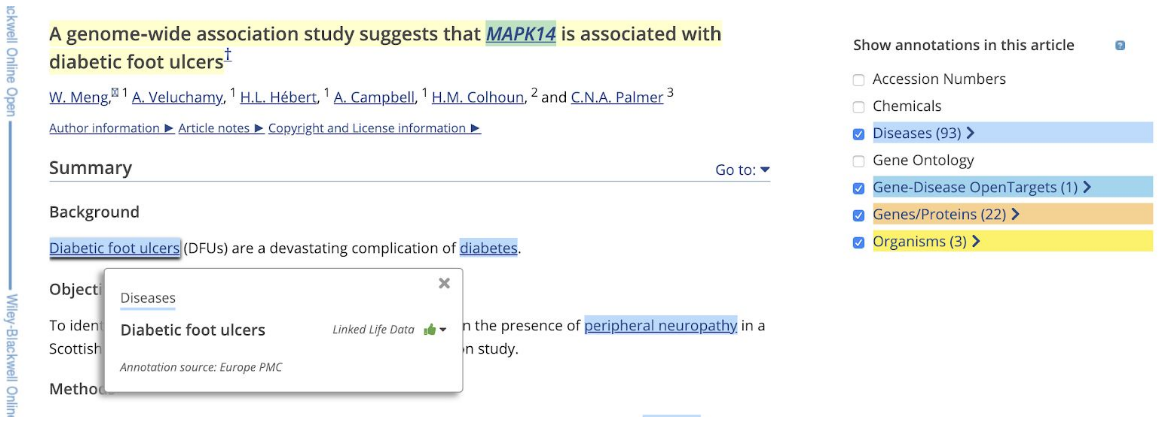
A genome-wide association study suggests that *MAPK14* is associated with diabetic foot ulcers[†]

W. Meng,[1] A. Veluchamy,[1] H.L. Hébert,[1] A. Campbell,[1] H.M. Colhoun,[2] and C.N.A. Palmer[3]

Author information ▶ Article notes ▶ Copyright and License information ▶

Summary Go to: ▾

Background

Diabetic foot ulcers (DFUs) are a devastating complication of diabetes.

Objecti[illegible]

To iden[illegible] n the presence of peripheral neuropathy in a Scottish [illegible] on study.

Metho[illegible]

Diseases

Diabetic foot ulcers — Linked Life Data

Annotation source: Europe PMC

Show annotations in this article

- Accession Numbers
- Chemicals
- Diseases (93)
- Gene Ontology
- Gene-Disease OpenTargets (1)
- Genes/Proteins (22)
- Organisms (3)

8. Use the mouse to select the entity that you would like to annotate.

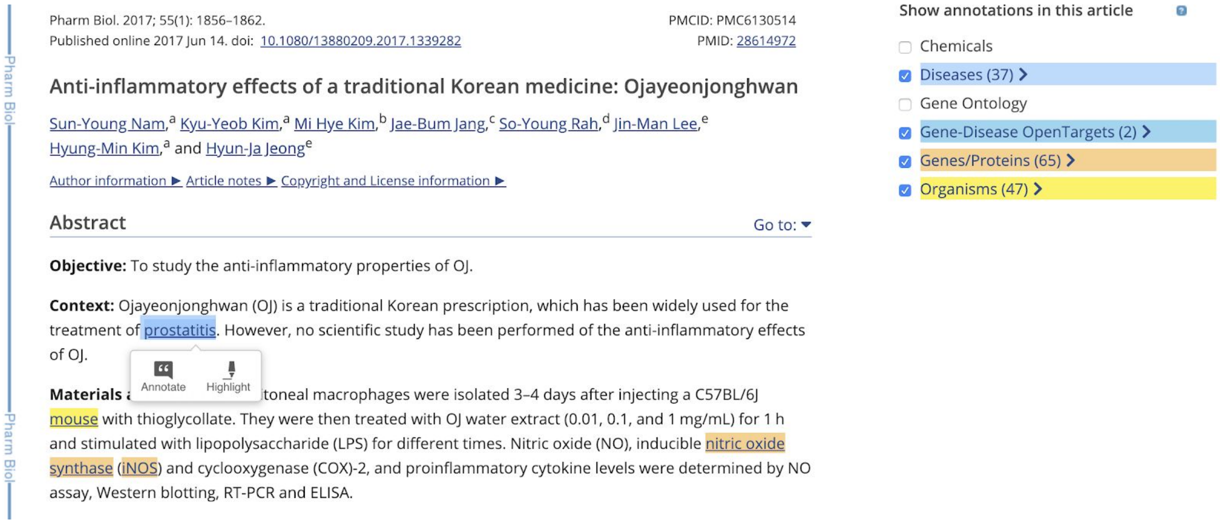
Pharm Biol. 2017; 55(1): 1856–1862.
Published online 2017 Jun 14. doi: 10.1080/13880209.2017.1339282

PMCID: PMC6130514
PMID: 28614972

Anti-inflammatory effects of a traditional Korean medicine: Ojayeonjonghwan

Sun-Young Nam,[a] Kyu-Yeob Kim,[a] Mi Hye Kim,[b] Jae-Bum Jang,[c] So-Young Rah,[d] Jin-Man Lee,[e] Hyung-Min Kim,[a] and Hyun-Ja Jeong[e]

Author information ▶ Article notes ▶ Copyright and License information ▶

Abstract Go to: ▾

**Objective:** To study the anti-inflammatory properties of OJ.

**Context:** Ojayeonjonghwan (OJ) is a traditional Korean prescription, which has been widely used for the treatment of prostatitis. However, no scientific study has been performed of the anti-inflammatory effects of OJ.

**Materials a**[illegible] toneal macrophages were isolated 3–4 days after injecting a C57BL/6J mouse with thioglycollate. They were then treated with OJ water extract (0.01, 0.1, and 1 mg/mL) for 1 h and stimulated with lipopolysaccharide (LPS) for different times. Nitric oxide (NO), inducible nitric oxide synthase (iNOS) and cyclooxygenase (COX)-2, and proinflammatory cytokine levels were determined by NO assay, Western blotting, RT-PCR and ELISA.

Annotate Highlight

Show annotations in this article

- Chemicals
- Diseases (37)
- Gene Ontology
- Gene-Disease OpenTargets (2)
- Genes/Proteins (65)
- Organisms (47)

9. Click “Annotate” to annotate the select words in the pop-up panel. Add tags of the annotation in the tag box according to the Tag Scheme. If you have any comments, you can leave it in the text box.

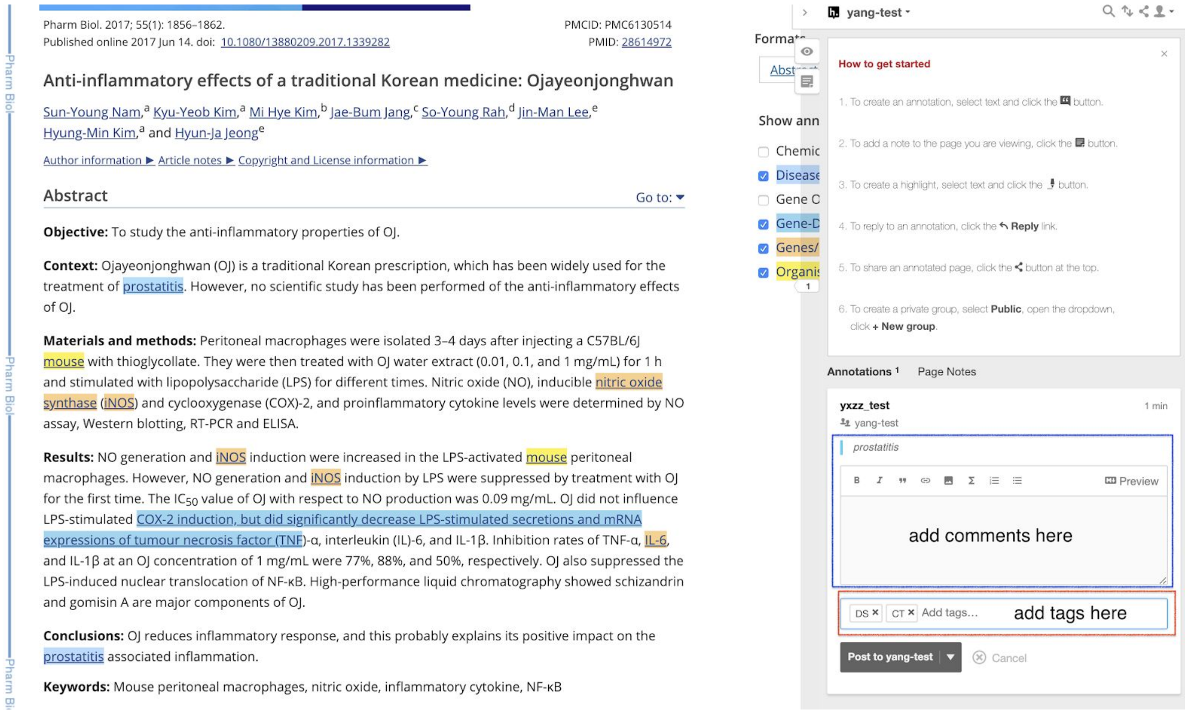

10. To finish the annotation, click the “Post to” button to post the annotation to the correct annotation group. Then the annotation will be added to the annotation group.

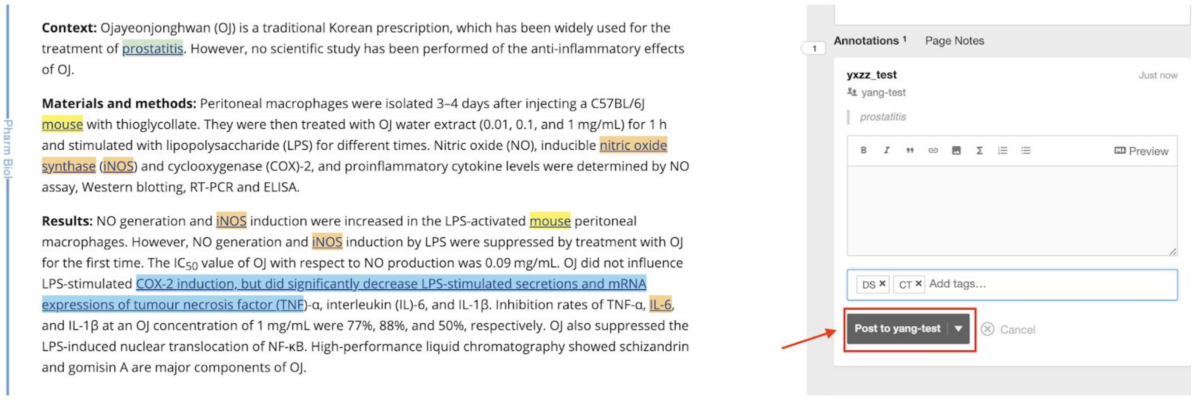

11. For Gene-Disease relationship annotation, click the highlighted text, the pre-annotated relationships will appear in a pop-up window.

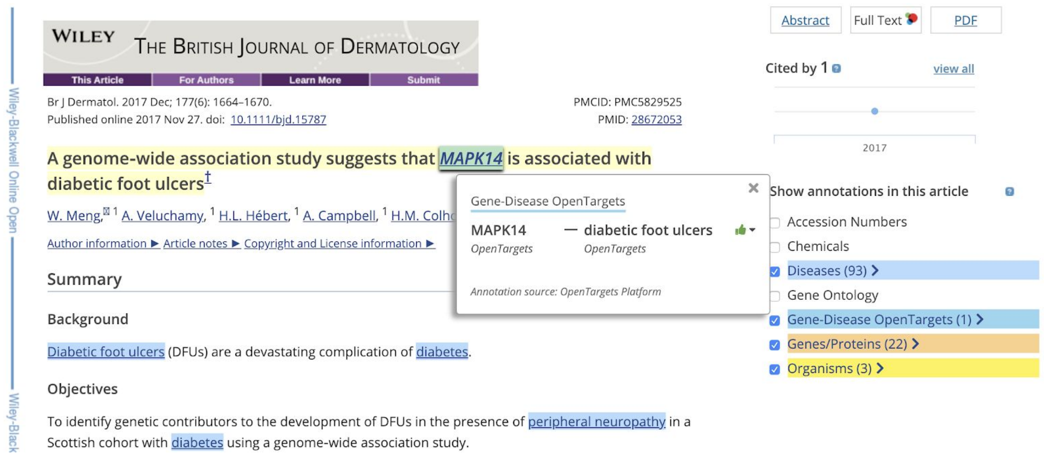

12. If a relationship between a gene and disease appears in the sentence, only select the part that contains the two entities using Hypothes.is. Then annotate the selected part by adding a gene-relationship tag to indicate whether it's a correct pre-annotation or a missing relationship annotation.

If the pre-annotation is wrong, select the pre-annotation and annotate it as a wrong relation.

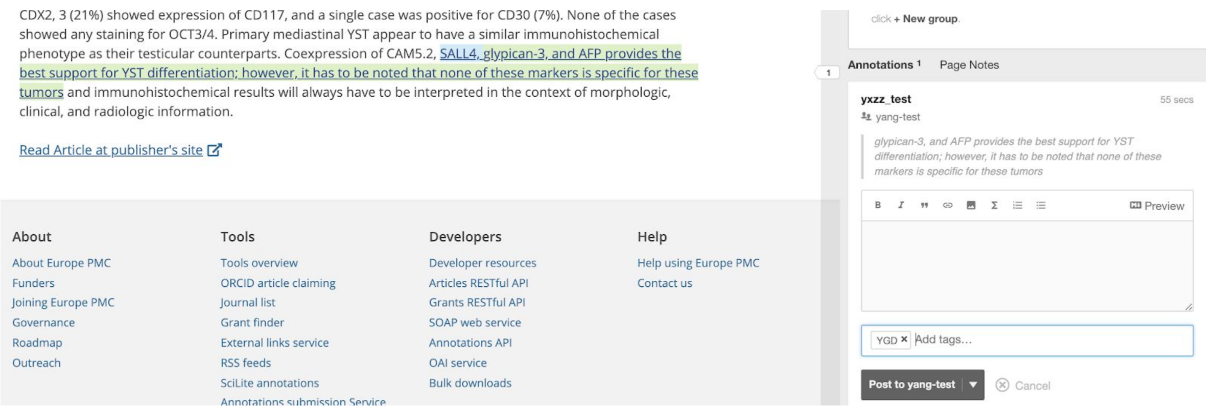
