## Supplementary material for "Europe PMC Annotated Full-text Corpus for Gene/Proteins, Diseases and Organisms": annotation guideline

Europe PMC Annotation Guidelines
Entity/Relationship Schema
Entity types
Gene/Protein-Disease relationship:
Boundary for selection of text
Entity annotations
Biomedical concepts
Annotate both singular or plural forms
Entities come after determiners “this, that, their, the, a, an, all, some, etc.”
Entity with hyphen
Entity with superscript, subscript and signs
Determine the span of annotations
Concepts within program or affiliation names
Concepts that are class/family names
Concepts that are composites of both the gene/protein and the source of organism
Concepts that are strain names
When a term is to be considered as a broad term
Validate pre-annotated annotations from EuropePMC
Relationship annotations

### Europe PMC Annotation Guidelines

For each identified entity and relationship, the selection of text must be semantically as close as possible to the concept and relationship described. The type the entity and relationship must be denoted using the guideline. These guides serve as a reference for consistently creating the annotations.

#### 1. Entity/Relationship Schema

##### 1. Entity types

If any text (word or phrase) is relevant to one of the following entity types, state the type of the selected text.

- Gene/Protein: Very broad terms like “DNA”, “RNA”, “gene”, etc should not be annotated. For uncertain terms, refer to Uniprot and Protein Ontology.
- Disease: For uncertain terms, refer to ULMS and EFO disease.

- Organism: Generic terms like “animal”, “human” are considered for annotation.

2. Gene/Protein-Disease relationship:

   If Gene/Protein and Disease entities are identified in the same sentence, check whether there exists a relationship between Gene/Protein entities and Disease entities. Select the relevant part of sentence if a relationship appears in the sentence and indicate the Gene/Protein and Disease entity pairs that are related.

   - Has relationship: The Gene/Protein and Disease entity pair has positive or negative association.
   - No relationship: The Gene/Protein and Disease entity pair has no association.

   If the relationship is ambiguous, annotators can mark the relationship annotation as “AMB” denoting “ambiguous”.

#### 2. Boundary for selection of text

For each entity annotation, any selected text span must be in the same sentence, i.e. the entity annotation must not start in current sentence and ends in the next sentence.

For each relationship annotation, the Gene/Protein and Disease entities involved in the relationship must be in the same sentence. i.e. in a relationship,  Gene/Protein entity appears in current sentence and Disease entity appears in the next sentence or vice versa.

#### 3. Entity annotations

To create an entity annotation, select a set of consecutive words in the documents that refers to entity types. For any give word or phrase, only annotate text that belongs to one of the entity types.

**NOTE: Examples are for illustrative purposes only and specific to each case, hence not all the entities are shown and highlighted.**

**RED: Gene/Protein BLUE: Disease GREEN: Organism**

##### a. Biomedical concepts

**Gene/Protein**: Annotations could be specific gene/protein names or classes/family names of gene/proteins. In particular, very broad concepts like “protein”, “gene”, “enzyme”, “receptors”, “kinase”, “cytokine”, “transcription regulators/factors” are out of the scope of annotations. However, family/subtype names of those concepts are considered for the annotations, such as “amylolytic enzyme”, “antioxidant enzyme”, “map kinase p38”, because these terms narrow the concepts to specific families of gene/protein, enzyme.

Annotators can refer to Uniprot and Protein Ontology.

**Disease**: Annotations could be specific disease names or classes/families of diseases. For example, “prostate tumor” and “tumor” are both valid concepts of disease. If “tumor” appears within a valid disease concept, e.g. “prostate tumor”, then that valid concept should be annotated as one entity.

**Organism**: Annotations could be specific species of organisms orclasses/families of species. For example, “mouse” and “animal” are both valid concepts of organism although “animal” is a very generic concept. Moreover, taxonomy families names are also considered for annotations, such as “asteraceae”, “cucurbitaceae” and “Lamiaceae”.

##### b. Annotate both singular or plural forms

The identified entity (including abbreviations) can be either singular or plural form as long as the entity is a valid concept of disease, organism or gene/protein.

Example 2.1:
Large **tumors** that have metastasized have a poorer prognosis than **tumors** that are confined to the breast. [PMC1885450]

In example 2.1, if the entity “**tumor**” or “**tumors**” has been annotated by the EuropePMC platform as **DISEASE**, it should be tagged as correct disease entity. Otherwise, it should be annotated as wrong disease entity. However, if it is not annotated by the platform, annotators don’t need to annotate it.

Example 2.2:
Finally, the researchers report that injection of **PKHB1** reduced the **tumor** burden in a **mouse** model of **CLL**. [PMC4348493]

Example 2.3:
Mild to moderate **bronchiolitis** and **pneumonia** were observed in the lungs of infected **animals**. [PMC2438613]

Example 2.4:
**HBV**, which is transmitted through contact with the blood or other bodily fluids of an infected person , can cause both acute (short-term) and chronic (long-term) **liver infections**. [PMC4280122]

Example 2.5:

Deletion of **SPDEF** in transgenic **mice** and cultures **prostate tumor** cells increased expression of **Foxm1** and its target genes.[PMC4177813]

Example 2.6:
**Pigs** also had a higher number of embedded **sand fleas** than all other species combined (p<0.0001). [PMC4608570]

##### c. Entities come after determiners “this, that, their, the, a, an, all, some, etc.”

Very often, there is a determiner (e.g. the, a, an, this, these, its, etc.) or quantifier (e.g. a lot of, some, most, each, several etc.) before an entity. In particular, numbers are used to give the information of quantity (e.g. ten tumors, 5 animals, etc.). Such words should **NOT** be included in the entity name as they are not biomedical concepts.

Example 3.1:
Sequencing of **KEAP1** in 12 cell lines and 54 **non-small-cell lung cancer** (**NSCLC**) samples revealed somatic mutations in **KEAP1** in a total of six cell lines and ten **tumors** at a frequency of 50% and 19%, respectively. [PMC1584412]

In the example 3.1, numbers such as “54” and “ten” are ignored as they are quantifiers and  not part of the biomedical terms.

Example 3.2:
None of the **lymphomas** in this group stained for the **LCV** viral capsid antigen (VCA) lytic marker. [PMC3464224]

In example 3.2, following the rules, “None of the” is not annotated.

Example 3.3:
This is an important concept since essentially all **humans** have life - long chronic **infections** from various **herpesviruses**. [PMC4298697]

In example 3.3, “all” is a quantifier and is therefore not included in the annotation.

Example 3.4:
We evaluated 18 **animals** with **malignancies** (16 **lymphomas**, one **fibrosarcoma** and one **carcinoma**) and 32 controls. [PMC6042791]

##### d. Entity with hyphen

In certain entity types, a hyphen may appear in the entity name e.g. in abbreviations. Hence, if the terms connected by the hyphen is a valid biomedical concept of gene/protein, disease or organism, it should be annotated as one entity. Otherwise, the terms on the left and right sides of a hyphen should be considered separately.

Example 4.1:
Pre - ART increases in Th17 and Th2 responses (e.g ., **IL-17**, **IL-4**) and lack of proinflammatory cytokine responses (e.g ., **G-CSF**, **GM-CSF**, **VEGF**) predispose individuals to subsequent **IRIS** , perhaps as biomarkers of immune dysfunction and poor initial clearance of CRAG. [PMC3014618]

In example 4.1, "IL-17" and "IL-4" are Gene/Protein names and therefore they annotated as shown in the example. In addition, separate "IL-17" into "IL" and "17" makes "17" senseless.

Example 4.2:
DRG axons began extending towards the localized **NT-3** source by the end of the first day and consistently displayed a strong chemoattraction by 3d in vitro, whereas they did not show such preference for **BSA**-loaded control beads (Figure 5A and 5B). [PMC529315]

In example 4.2, "NT-3" is the abbreviation of Gene/Protein name of Neurotrophin-3 and thus annotated as Gene/protein entity. However, "BSA-loaded control beads" is not a biomedical concept of Gene/Protein, disease and organism. In this case, only "BSA" on the left side of the hyphen is annotated as the gene/protein entity.

Example 4.3:
Small genetic contributions could also be seen from the susceptibility genes of RA identified so far, including **HLA-DR4**, **PADI4**, **PTPN22** and **FCRL3** [6-9]. [PMC1860061]

In example 4.3, "HLA-DR4" together is a Gene/Protein name and therefore is annotated as one Gene/Protein entity.

Example 4.4:
Because **VEGF** is a key regulator of **tumor** development , several anti-**VEGF** therapies drugs that target **VEGF** and its receptors have been developed.

In example 4.4, "VEGF" should be annotated instead of "anti-VEGF" because "anti-VEGF therapies drugs" is not a biomedical concept of gene/protein, disease and organism. Thus, we only annotate "VEGF" which is a concept listed in this guideline.

##### e. Entity with superscript, subscript and signs

Superscripts and subscripts are irrelevant to biomedical concepts and should **NOT** be included in annotations.

Example 5.1
(H) Fibroblast-like cells present in the bone shaft of **Bmp2**$^{C/C}$; **Bmp4**$^{C/C}$; Prx1::cre mouse. [PMC1713256]

In example 5.1, the superscript $^{C/C}$ is not part of the concept and should not be included in the annotation.

Example 5.2:
**Stat5a** is suggested to contribute to tolerance through maintenance of the **CD4**+**CD25**+ regulatory T cell population [35].

In example 5.2, signs like "+" should not be annotated as it usually is not a part of a concept.

Example 5.3:
Since < 1% of **Trip13**$^{Gt/Gt}$ pachytene nuclei had normal repair (as judged by absence of persistent DSB repair markers ; see above), but most of the pachytene nuclei had **MLH1**/3 foci , it was unlikely that the **MLH1**/3 foci formed only on chromosomes with fully repaired DSBs. [PMC1941754]

In example 5.3, following the guideline, superscript $^{Gt/Gt}$ is not annotated as part of the concept.

Example 5.4:
When we compared the aggregation curves of human platelets from a healthy donor with the ones obtained from an individual with a **von Willebrand factor type 1 defect** , we found that the difference in the curves was much more pronounced as observed in our studies of healthy mouse platelets and **anxA7**$^{-/-}$ platelets. [PMC194730]

In example 5.4, following the guideline, superscript $^{-/-}$ is not annotated as part of the concept.

#### f. Determine the span of annotations

Sometimes, a potential concept can be a complex noun phrase. Thus, it's important to determine the right span of the annotation to make valid annotations.

The basic principle and procedure to determine the right span is,

(1) follow the previous steps a, b, c, d and e first to ignore quantifiers, determiners, superscript, etc.
(2) if the phrase is a valid concept of gene/protein, disease or organism, then annotate it as one of the concepts.

(3) if the phrase is not related to any concept, you should try to find any valid concepts within the phrase i.e. only part of the phrase is annotated.

Example 6.1:
Encouraged by the promising clinical activity of **epidermal growth factor receptor** (**EGFR**) kinase inhibitors in treating **glioblastoma** in **humans**, we have sequenced the complete **EGFR** coding sequence in **glioma tumor** samples and cell lines. [PMC1702556]

In example 6.1, "glioblastoma in humans" is a phrase but "glioblastoma" and "humans" should be individually annotated because "in" is a preposition and should not be included in the concept annotation. "the complete EGFR coding sequence" is a phrase but it is not related to any concept in the guideline, hence, within the phrase, "EGFR" is a valid gene/protein concept and should be annotated.

Example 6.2:
Katharina Kranzer and colleagues investigate the operational characteristics of an active **tuberculosis** case-finding service linked to a mobile **HIV** testing unit that operates in underserviced areas in Cape Town, South Africa. [PMC3413719]

In example 6.2, "HIV" is annotated instead of "a mobile HIV testing unit" because a testing unit is not a biomedical concept. Similarly, the phrase "active tuberculosis case-finding service" is not a valid biomedical concept and therefore only "tuberculosis" is annotated as a valid disease concept.

Example 6.3:
**Severe acute respiratory syndrome** (**SARS**) is a **flu**-like illness and was first recognized in China in 2002, after which the disease rapidly spread around the world.

In example 6.3, "flu-like illness" is not a valid biomedical concept and therefore only "flu" is annotated as a disease concept.

Example 6.4:
Two recent papers provide new evidence relevant to the role of the **breast cancer** susceptibility gene **BRCA2** in DNA repair. [PMC138691]

In example 6.4, "breast cancer susceptibility gene" is describing/explaining "BRCA2" and it is not a specific gene name. Therefore, it should not be annotated as one entity. Instead, within the phrase, "breast cancer" should be annotated.

Example 6.5:

When we compared the aggregation curves of human platelets from a healthy donor with the ones obtained from an individual with a **von Willebrand factor type 1 defect** , we found that the difference in the curves was much more pronounced as observed in our studies of healthy mouse platelets and **anxA7**$^{-/-}$ platelets. [PMC194730]

In example 6.5, “von Willebrand factor type 1 defect” should be annotated as one entity because together it is a valid disease name, which is the ” type 1 defect” of the gene/protein “von Willebrand factor”.

Example 6.6:
Whole mount immunohistochemical analysis of embryos using a **CD31** antibody as described. [PMC324396]

In example 6.6, although “CD31” describes “antibody”, “antibody” should not be annotated because “CD31” is the main concept in this phrase. (better explanation required)

Example 6.7:
**Human** infective **Trypanosoma brucei rhodesiense** were detected in 21.5% of **animals** infected with **T. brucei s.l.** [PMC3022529]

In example 6.7, in the phrase “animals infected with T. brucei”, “animals” and “T. brucei” should be annotated separately because the longer form is not an organism name. The same reason for breaking “Human infective Trypanosoma brucei rhodesiense” into two separate annotations.

Example 6.8:
Earlier initiation of antiretroviral therapy may be a key component of global and national strategies to control the **HIV**-associated **tuberculosis** syndemic. [PMC3404110]

In example 6.8, the phrase, “HIV-associated tuberculosis syndemic” is not a biomedical concept of either organism, disease and gene/protein. Therefore, we only annotate “HIV” and “tuberculosis”.

#### g. Concepts within program or affiliation names

Some valid concepts may appear in affiliation names, however they should not be annotated as semantically they are not part of the research.

Example 7.1:
Cancer Research UK provides information on all aspects of **brain tumors** for patients and their caregivers. [PMC2621261]

Example 7.2:

US National Cancer Institute information for patients and professionals on **lung cancer** (in English and Spanish). [PMC2043012]

Example 7.3:
An overview of **HIV infection** and **AIDS** is available from the US National Institute of Allergy and Infectious Diseases.

In example 7.1, 7.2 and 7.3, the concepts, for example "Cancer" and "Allergy" are not annotated because they are part of the affiliation names.

##### h. Concepts that are class/family names

Class/family names are also considered for annotations, such as "asteraceae", "cucurbitaceae" and "Lamiaceae".

Example 8.1:
**Cucurbitaceae** represent an important plant family in which many species contain cucurbitacins as secondary metabolites synthesized through isoprenoid and triterpenoid pathways.

##### i. Concepts that are composites of both the gene/protein and the source of organism

In some cases, the concept is a composite of both gene/protein and the source of organism, such as "CsbHLH18", which should be annotated as Gene/Protein.

Example 9.1:
The transcription factor **CsbHLH18** of sweet orange functions in modulation of cold tolerance and homeostasis of reactive oxygen species by regulating the antioxidant gene.

##### j. Concepts that are strain names

In the case that the strain of an organism is mentioned along with the organism name, the strain name should be annotated. If the strain name is mentioned standalone without organism name, it is not considered for annotations.

Example 10.1
Here we show that the addition of FOS to ***P. aeruginosa*** **PAO1** cultures decreases growth and biofilm formation.

Example 10.2
In order to test this hypothesis, we infected rat primary monocyte cultures with PAO1 and measured cytokine release in the presence and absence of oligosaccharides.

In the example 10.1, the strain name “PAO1” is mentioned with the organism name “P. aeruginosa”. As such “P. aeruginosa PAO1” should be annotation as one ORGANISM concept. However, in example 10.2, only “PAO1” is mentioned and therefore it should not be considered for annotation.

##### k. When a term is to be considered as a broad term

In general, very broad terms are not useful and hence should not be considered for annotation. Examples of very broad terms are “gene”, “protein”, “enzyme”, “receptor” and their plural forms. However, as mentioned in section **3.h**, class/family names are not considered as very broad terms when they represent specific groups of concepts. In addition to section **3.h**, when a very broad term is described by adjectives, etc. that make the concept more specific, they should be annotated as one concept.

Some examples of terms that are considered for annotations are : transcription regulator, transcription factor, phosphoproteins, kinase, antioxidant enzyme, cytokine, tyrosine kinase, receptor tyrosine kinase, etc.

However, there are some special cases to look at:
“liver infection” vs “pig infection” vs “bacterial infection”

“pig infection” is not a disease concept because pig is the species that got infected.

“bacterial infection” is a disease concept because the bacterial leads to the infection. Similar valid concepts are “virus infection”, “HIV infection”, etc.

“Liver infection” is a disease concept because the liver is the exact location that infection occurs. Similar valid concepts are “lung infection”, “ear infection”, etc.

##### l. Validate pre-annotated annotations from EuropePMC

Existing EuropePMC annotations may cover very generic terms such as “infection” and “acute illness” but as long as the annotation is correct (e.g. it is not part of an organisation name like “*animal* protection organization” or wrong type/span), it should be annotated as correct. However, such very generic terms DO NOT need to be annotated by annotators if they are missing.

#### 4. Relationship annotations

To create a Gene-Disease relationship annotation, select sentences in the documents that:

- contain entities of both gene and disease
- have a relationship between gene and disease entities.

A relationship indicates association of gene and disease entities, either positive or negative associations. For given documents, only annotate the part of sentences that have gene-disease relationships. If a gene-disease relationship exists, then the relationship and the gene-disease entities that establish the relationship should be annotated explicitly.

In the following examples, gene and disease entities are annotated and the relationships are listed explicitly.

a. Positive association

   A relationship with positive association indicates that one entity influences the other one. No matter if the influence is positive or negative.

   Example 8.1:
   Specific hypermethylation of **NEUROG1** and **NR2E1** was identified as a feature of **cortical tumours**. [PMC6068350]

   Gene-disease relationships:
   NEUROG1 - cortical tumors
   NR2E1 - cortical tumors

   Example 8.2:
   **Human epidermal growth factor receptor 2** (***Erbb2***/**HER2**) overexpression, which was previously detected in invasive **breast cancer**, has now been implicated in advanced **gastric cancer** (**GC**) and **gastroesophageal junction cancer** (**GEC**). [PMC5948243]

   Gene-disease relationships:
   Human epidermal growth factor receptor 2 - breast cancer
   Erbb2 - breast cancer
   HER2 - breast cancer

   Human epidermal growth factor receptor 2 - gastric cancer
   Erbb2 - gastric cancer
   HER2 - gastric cancer

   Human epidermal growth factor receptor 2 - GC
   Erbb2 - GC
   HER2 - GC

   Human epidermal growth factor receptor 2 - gastroesophageal junction cancer
   Erbb2r - gastroesophageal junction cancer
   HER2 - gastroesophageal junction cancer

Human epidermal growth factor receptor 2 - GEC
Erbb2r - GEC
HER2 - GEC

Example 8.3:
**HER2** overexpression was significantly more common in diffuse type than in intestinal type of **tumors** (39.8 vs. 14.9%; $p < 0.001$). [PMC5948243]

Gene-disease relationships:
HER2 - tumors

Example 8.4:
**HER2** overexpression was evident in nearly 25% of the Malaysian patients with locally advanced or metastatic **gastric cancer**. [PMC5948243]

Gene-disease relationships:
HER2 - gastric cancer

Example 8.5:
The therapeutic index of **rheumatoid arthritis** (**RA**) may be improved with MTX therapy based on the **IL-6** circadian rhythm. [PMC5884908]

Gene-disease relationships:
IL-6 - rheumatoid arthritis
IL-6 - RA

Example 8.6:
Despite similar demographics, co-morbidities, valve narrowing, **myocardial hypertrophy**, and **fibrosis**, patients with asymmetric wall thickening had increased **cardiac troponin I** and brain natriuretic peptide concentrations (both $P < 0.001$).
[PMC5837366]

Gene-disease relationships:
Cardiac troponin I - myocardial hypertrophy
Cardiac troponin I - fibrosis

Example 8.7:
Increased expression of the **TRPM4** channel has been reported to be associated with the progression of **prostate cancer**. [PMC5792731]

Gene-disease relationships:
TRPM4 - prostate cancer

Example 8.8:

**TRPM4** expression is increased in the transition from prostatic intraepithelial neoplasia (PIN) to **prostate cancer** (Ashida *et al*., 2004; Singh *et al*., 2006). [PMC5792731]

Gene-disease relationships:
TRPM4 - prostate cancer

Example 8.9:
**Akt1** activation is regulated by Ca2+/**CaM** and **TRPM4** in **prostate cancer** cells. [PMC5792731]

Gene-disease relationships:
Akt1 - prostate cancer
CaM - prostate cancer
TRPM4 - prostate cancer

Example 8.10:
On the other hand, deregulation of **Akt** signaling is a common alteration in **prostate cancer** (Li *et al*., 2005). [PMC5792731]

Gene-disease relationships:
Akt - prostate cancer

Example 8.11:
Several studies on **prostate cancer** have suggested that the expression of **TRPM4** is a relevant event in the progression of this **tumor** (Holzmann *et al*., 2015; Schinke *et al*., 2014). [PMC5792731]

Gene-disease relationships:
TRPM4 - prostate cancer
TRPM4 - tumor

Example 8.12:
Importantly, the analysis of 10 gene expression datasets from patients with **prostate cancer** and their controls shows that the most enriched pathway coexpressed with the **TRPM4** gene is the **Wnt** signaling pathway, supporting our *in vitro* results and sustaining a relationship between the expression of this channel and the activity of this signaling pathway in **prostate cancer** (Fig. S5). [PMC5792731]

Gene-disease relationships:
TRPM4 - prostate cancer
Wnt - prostate cancer

Example 8.13:

Serum **tissue factor** as a biomarker for **renal clear cell carcinoma** [PMC5815530]

Gene-disease relationships:
tissue factor - renal clear cell carcinoma
A relationship exist as “biomarker for” indicates a relationship.

Example 8.14:
Genetic variants in five genes (***MIA3***, ***MRAS***, ***P2RX7***, ***CAMKK2***, and ***SMAD3***) were associated with increased waist circumference in patients with **schizophrenia** spectrum disorder ($P$<0.046). [PMC5662154]

Gene-disease relationships:
MIA3 - schizophrenia
MRAS - schizophrenia
P2RX7 - schizophrenia
CAMKK2 - schizophrenia
SMAD3 - schizophrenia

Example 8.15:
Genetic variants in the ***PPARD***, ***MNTR1B***, ***NOTCH2***, and ***HNF1B*** were nominally associated with **schizophrenia** spectrum disorder irrespective of waist circumference ($P$<0.027). [PMC5662154]

Gene-disease relationships:
PPARD - schizophrenia
MNTR1B - schizophrenia
NOTCH2 - schizophrenia
HNF11B - schizophrenia

Example 8.16:
The reported risk alleles of genetic variants rs10830963 in ***MTNR1B*** and rs10923931 in ***NOTCH2*** were associated with **diabetes mellitus type 2**-related traits in GWA studies ($P<5\times10^{-8}$) (Zeggini *et al.*, 2008; Prokopenko *et al.*, 2009). [PMC5662154]

Gene-disease relationships:
MTNR1B - diabetes mellitus type 2
NOTCH2 - diabetes mellitus type 2

Example 8.17:
Heterozygous mutations in *UMOD* encoding the urinary protein **uromodulin** are the most common genetic cause of **autosomal dominant tubulointerstitial kidney disease** (**ADTKD**). [PMC5837645]

Gene-disease relationships:

uromodulin - autosomal dominant tubulointerstitial kidney disease
uromodulin - ADTKD

Example 8.18:
Curcumin effectively protected mice from **sepsis** as evidenced by decreasing histological damage, reducing **AST** (352.0 vs 279.3 U/L), **BUN** (14.8 vs 10.8 mmol/L) levels and the proportion of macrophages in spleen (31.1% vs 13.5%). [PMC6130682]

Gene-disease relationships:
AST - sepsis
BUN - sepsis

Example 8.19:
These results suggest that isotalatazidine hydrate is a potent dual **cholinesterase** inhibitor and can be used as a target drug in **Alzheimer diseases**. [PMC6130761]

Gene-disease relationships:
Cholinesterase - Alzheimer diseases

Example 8.20:
A genome-wide association study suggests that **MAPK14** is associated with **diabetic foot ulcers**. [PMC5829525]

Gene-disease relationships:
MAPK14 - diabetic foot ulcers

Example 8.21:
In humans, low **serum carnosinase** (**CN1**) activity protects patients with **type 2 diabetes** from **diabetic nephropathy**. [PMC6009930]

Gene-disease relationships:
serum carnosinase - diabetic nephropathy
CN1 - diabetic nephropathy
serum carnosinase - type 2 diabetes

Example 8.22:
Cysteine-compounds influence the dynamic behaviour of **CN1** and therefore present a promising option for the treatment of **diabetes**.

Gene-disease relationships:
CN1 - diabetes

b. Negative association

   A relationship with negative association indicates that there doesn't have influence between one entity and the other.

   Example 8.23:

   Despite an amplified biological effect of the homozygote mutation, the proband did not show a strikingly more severe clinical evolution nor was the near absence of urinary **uromodulin** associated with **urinary tract infections** or **kidney stones**.[PMC5837645]

   Gene-disease relationships:
   uromodulin - kidney stones
   uromodulin - urinary tract infections

   Example 8.24:

   There was no statistically significant correlation between **HER2** positivity and patient age, race, tumor location, **tumor** differentiation, and TNM staging.[PMC5948243]

   Gene-disease relationships:
   HER2 - tumor

c. No association

   No association indicates that no relationship between one entity and the other. It occurs sometimes in literature that gene and disease entities are mentioned in the sentence but not mentioning any association.
